# BRD8 Disruption Unlocks Retrotransposon-Driven Tumor-Intrinsic Immunogenicity via Redistribution of Histone Acetylation in Liver Cancer

**DOI:** 10.64898/2026.01.21.700714

**Authors:** Ruijie Gong, Yijie Shen, Jianhang Huang, Jie Wang, Siyuan Xu, Songchen Liu, Qin Feng, Jia Yu, Jianing Zhao, Jia Fan, Jiabin Cai, Xianjiang Lan

**Affiliations:** Department of Hepatobiliary Surgery and Liver Transplantation, Liver Cancer Institute, Zhongshan Hospital, Institutes of Biomedical Sciences, Key Laboratory of Carcinogenesis and Cancer Invasion of Ministry of Education, Key Laboratory of Medical Epigenetics and Metabolism, Zhongshan Hospital, Fudan University, Shanghai 200032, China; Department of Systems Biology for Medicine, School of Basic Medical Sciences, Liver Cancer Institute, Zhongshan Hospital, Fudan University, Shanghai 200032, China; Endoscopy Center and Endoscopy Research Institute, Zhongshan Hospital, Fudan University, Shanghai 20032, China; Department of Liver Surgery, Zhongshan Hospital (Xiamen Branch), Fudan University, Xiamen 361015, China

## Abstract

Overall response rate for immune checkpoint blockade (ICB) therapy remains limited in advanced hepatocellular carcinoma (HCC). Epigenetic perturbation has been shown to enhance tumor-intrinsic immunogenicity and thus reverse resistance to immunotherapy in multiple cancer types. Here, through domain-focused CRISPR/Cas9 genetic screens targeting chromatin regulators, we identified the bromodomain-containing protein 8 (BRD8) as a novel epigenetic factor that functions to suppress tumor-intrinsic immunity through BRD8/EP400 chromatin remodeling complex in HCC. Depletion of BRD8 potently induces anti-tumor immune responses and sensitizes tumors to ICB therapy in mouse tumor models. Mechanistically, BRD8 knockout leads to partial redistribution of H3K27ac to repetitive elements, activating double-stranded RNA (dsRNA) accumulation and subsequent type I interferon (IFN) response, leading to tumor rejection via remodeling the tumor microenvironment. The first-in-class selective chemical probe DN02 targeting the bromodomain of BRD8 also exhibits immunomodulatory and anti-tumor effects. In line with these findings, low BRD8 expression correlated with enriched interferon signatures, elevated immune cell infiltration, and prolonged survival in HCC patients. Together, this study reveals BRD8 as a selective epigenetic vulnerability for promoting tumor immunogenicity, and its bromodomain as a druggable target to overcome immunotherapy resistance in liver cancer patients.

**Significance:** The induction of viral mimicry by epigenetic interference could trigger tumor-intrinsic immune response and enhance immunotherapy efficacy. Here, through domain-focused CRISPR/Cas9 screening, we identify BRD8 complex as a regulator involved in the suppression of repetitive elements by controlling H3K27ac distribution and propose BRD8 as a novel target to enhance anti-tumor immunity.

## Introduction

Primary liver cancer was the sixth most common tumor and the third mortality malignant tumor worldwide in 2022(1). The number of new liver cancer and related mortality are predicted to reach 1.52 million and 1.37 million respectively in 2050 if current intervention is not improved(2). Hepatocellular carcinoma (HCC) is the most common pathological type of primary liver cancer(3), which a considerable number of patients were diagnosed at an advanced stage and cannot undergo curative surgery. Although comprehensive treatment based on PD-1/PD-L1 immune checkpoint blockade (ICB) therapy has shown certain benefit in some patients, the overall objective response rate (ORR) of these studies were less than 30% and outcomes remain dismal(4,5). This emphasizes the importance of improving immunotherapeutic efficacy for HCC patients. Interferon (IFN)-related signature genes, such as expression of PD-L1, accurately predicts response to ICB therapy(6,7). Therefore, there is an urgent clinical need to identify novel targets to enhance tumor-intrinsic IFN response and develop combination regimens that ultimately sensitize tumors to ICB therapy.

Triggering a tumor-intrinsic IFN response represents a pivotal strategy for converting immunologically cold tumors into immune-inflamed (hot) states, thereby sensitizing them to ICB therapy(8). Recently, epigenetic perturbations have converged with tumor-intrinsic IFN response through a phenomenon known as “viral mimicry”(9–13). Interfering with the transcriptional regulation of transposable elements (TEs)-which make up ∼50% of the human genome-through DNA methylation, histone modifications, and chromatin remodeling, can induce tumor cell-intrinsic accumulation of endogenous virus-mimicking nucleic acids, such as double-stranded RNA (dsRNA)(14). Recognition of these nucleic acids by innate immune sensors activates IFN signaling, mimicking an antiviral response. The resulting IFN-driven inflammation in tumor cells is highly immunogenic, and therapeutic strategies inducing this viral mimicry have shown significant preclinical(10,15,16) and clinical potential(17,18). However, the role of modulating the tumor-intrinsic IFN response specifically in HCC remains largely underexplored.

Using domain-focused CRISPR/Cas9 screening targeting chromatin regulators(19), we identified bromodomain-containing protein 8 (BRD8) as a critical epigenetic repressor of the tumor-intrinsic IFN response in HCC. BRD8, a subunit of the EP400 complex and a reader of acetylated lysine residues (KAc)(20), has been reported to be implicated in carcinogenesis(21,22), cell proliferation(23–26), metastasis(27) and immune regulation(26,28,29) across cancers. We demonstrate that BRD8 maintains histone H3 lysine 27 acetylation (H3K27ac) at its bound regions, and disruption of BRD8 redistributes H3K27ac to poised enhancers at TE loci, thereby derepressing TEs and driving dsRNA accumulation. Genetic BRD8 deletion or pharmacological bromodomain inhibition induces a potent cell-intrinsic IFN response in vitro and enhances tumor immunogenicity in vivo, significantly sensitizing HCC mouse models to ICB therapy. Our findings reveal a novel epigenetic mechanism linking H3K27ac redistribution to retroelement-driven dsRNA accumulation and establish BRD8-targeting as a promising strategy to overcome ICB resistance in liver cancer treatment.

## Result

### CRISPR/Cas9 screening identifies BRD8, EP400 and TIP60 as novel regulators of CD47 and PD-L1

CD47 and PD-L1 are the two important immune checkpoints and they were able to be induced by IFN-γ treatment in HCC (Supplementary Fig. S1a-c), thus we chose them as the markers of tumor-intrinsic interferon response activation, and attempted to identify key epigenetic factors that modulate their expression and/or the interferon signaling pathways. To this end, we performed CRISPR genetic screening using a library targeting 196 human chromatin regulator domains covering the main epigenetic writers, erasers and bromodomain proteins to search novel regulators of basal and IFN-γ-induced CD47 and PD-L1 expression in Huh7 cells (Fig. 1a). The CD47/PD-L1 high and low expression cells were purified by anti-CD47 or PD-L1 fluorescence activated cell sorting (FACS) and then subject to deep sequencing to assess the representation of each sgRNA in the two cell populations, respectively (Figure. 1b).

**Fig. 1.**
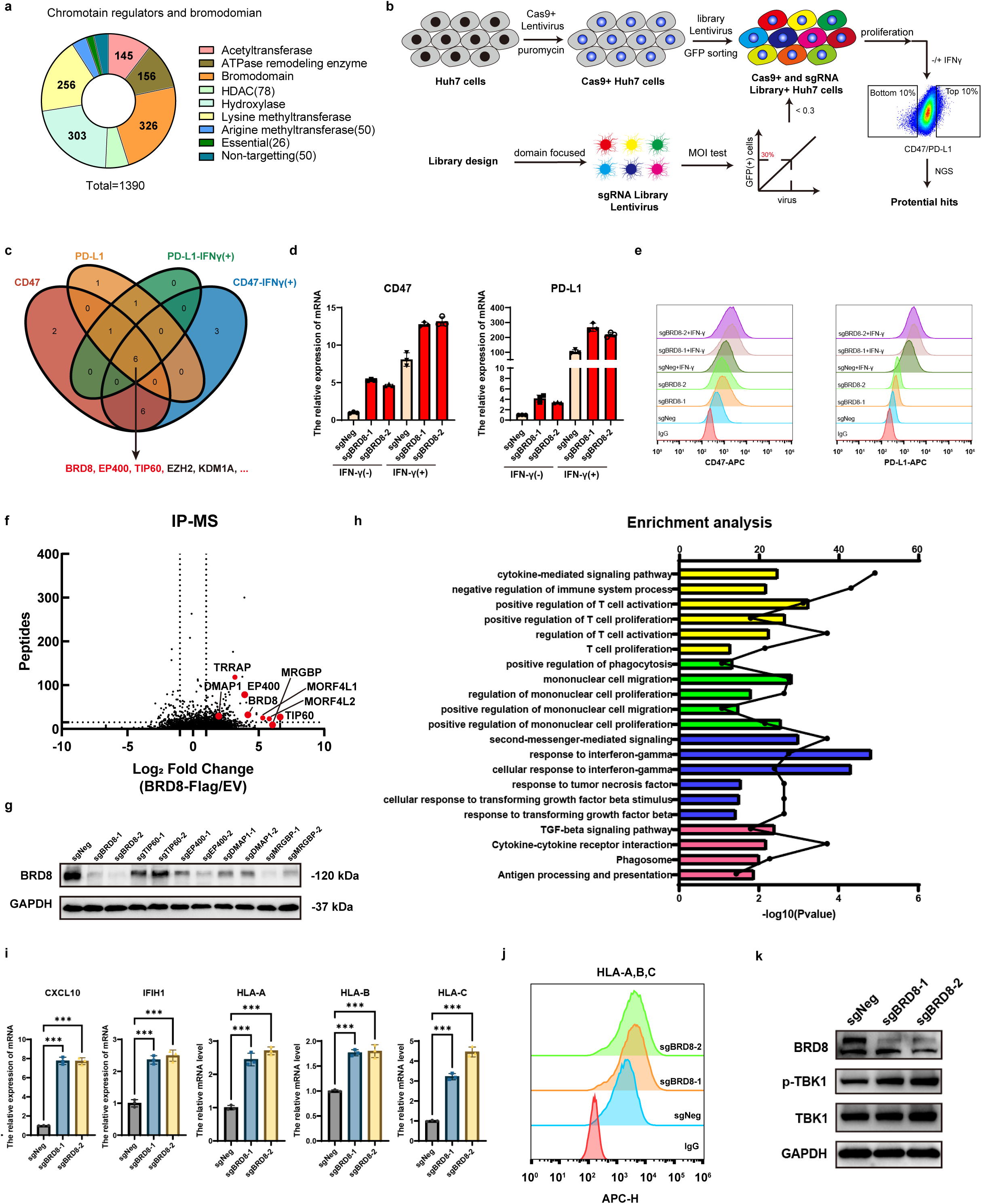
Domain-focused CRISPR screens identify BRD8 as a regulator of cell-intrinsic interferon response. (**a**) Composition of the CRSIPR library. (**b**) Schematic of the screening strategy. FACS, fluorescence-activated cell sorting; NGS, Next-generation sequencing. (**c**) Venn diagram of the four screen results. (**d**) CD47 or PD-L1 mRNA levels in Huh7 cells upon BRD8 depletion with or without IFNγ treatment for 48 h. Results is shown as mean ± SD (n = 3). (**e**) CD47 or PD-L1 flow cytometry analyses of WT and BRD8-depleted Huh7 cells with or without IFNγ treatment for 48 h. (**f**) Scatter plot depicting the enrichment fold-change of immunoprecipitated proteins using Flag-tagged BRD8 compared to empty vector control in IP-MS assays. (**g**) Western Blot of BRD8 in control (sgNeg) and EP400 complex partners knockout (BRD8, TIP60, EP400, DMAP1, MRGBP) in Huh7 cells. GAPDH was used as loading control. (**h**) Gene ontology (GO) and Kyoto Encyclopedia of Genes and Genomes (KEGG) analysis of RNA-seq data showing immune-related pathways that are up-regulated upon BRD8 depletion in Huh7 cells. (**i**) ISG, cytokine or MHC mRNA levels in Huh7 cells upon BRD8 depletion. Results is shown as mean ± SD (n = 3). (**j**) HLA-A, B, C flow cytometry analyses of WT and BRD8-depleted Huh7 cells. (**k**) Western Blot of total or phosphorylated TBK1 in control and BRD8 knockout in Huh7 cells. GAPDH was used as loading control.

As expected, all sgRNAs targeting EZH2 and KDM1A (also named LSD1), known negative regulators of tumor-intrinsic interferon response(11,16,30), were enriched in the CD47 and PD-L1-high populations (Fig. 1c and Supplementary Fig. S1d-g), validating our screens. Interestingly, the sgRNAs targeting BRD8, EP400 and TIP60, subunits of the EP400 complex, were also enriched in the two cell populations (Fig. 1c and Supplementary Fig. S1h-k), suggesting that they are negative regulators of CD47 and PD-L1. To further validated the screening results, we generated BRD8-knockout (KO) Huh7 cells with two independent sgRNAs (Supplementary Fig. S2a). Consistently, BRD8 depletion strongly elevated the mRNA and protein levels of CD47 and PD-L1 with or without IFN-γ treatment (Fig. 1d-e). We also obtained the similar results for EP400 and TIP60 in Huh7 cells (Supplementary Fig. S2b-c). These results demonstrate that BRD8, EP400 and TIP60 are novel repressors of CD47 and PD-L1.

### BRD8 represses CD47 and PD-L1 expression through the EP400 complex but independent of its acetyltransferase activity and deposition of H2A.Z

BRD8 is a well-known subunit of the EP400 chromatin remodeling complex(31). To further investigate whether BRD8 regulates CD47 and PD-L1 expression through the EP400 complex, we conducted Flag-BRD8 immunoprecipitation–mass spectrometry (IP–MS) experiments in Huh7 cells. Proteomics results showed that BRD8 was mainly associated with components of the EP400 chromatin remodeling complex, including EP400, TIP60, and additional subunits TRRAP, DMAP1, MRGBP, MORF4L1, MORF4L2 (Fig. 1f and Supplementary Fig. S2g). Notably, depletion of the other subunit DMAP1 or MRGBP also substantially induced the expression of CD47 and PD-L1(Supplementary Fig. S2e-f). Furthermore, to interrogate the functional relationship between BRD8 and other components of the EP400 complex, we found that depletion of EP400, TIP60, DMAP1 or MRGBP diminished the protein abundance of BRD8 to varying degrees (Fig. 1g), consistent with a prior study(32). Collectively, these findings demonstrate that BRD8 functions as a repressor of CD47 and PD-L1 through the EP400 complex. In line with the results in Huh7 cells, BRD8 deficiency in another two human HCC cell line HepG2 and MHCC-97L also led to the upregulation of CD47 and PD-L1 (Supplementary Fig. S2g-l), confirming the general role of BRD8 in HCC cell lines. Considering the fact that BRD8 ranks top in the CRISPR screens (Top 4 in the CD47 screen and Top 2 in the PD-L1 screen), we focused on BRD8 in subsequent studies.

The BRD8/EP400 complex influences chromatin structure through two enzymatic activities(33–35): histone acetylation and ATP-dependent exchange of histone variant H2A.Z (encoded by H2AFZ and H2AFV)(36,37). We first asked whether BRD8/EP400 complex regulating CD47 and PD-L1 relies on its acetyltransferase activity. Notably, treatment of TIP60 enzymatic activity inhibitors NU9056 or MG149 failed to affect the expression of CD47 and PD-L1 in Huh7 cells (Supplementary Fig. S3a-d), suggesting that the acetyltransferase activity of BRD8/EP400 complex is disposable for the regulation of CD47 and PD-L1 expression. Moreover, H2A.Z deposition has been reported to be maintained by the BRD8/EP400 complex and involved in the transcriptional repression in promoter region of target genes(32,38,39). Indeed, BRD8 depletion led to marked loss of H2A.Z occupancy at genome-wide regions including the CD47 and PD-L1 promoter without affecting H2A.Z protein abundance (Supplementary Fig. S3e-g). However, no increased expression of CD47 and PD-L1 were observed in H2A.Z-deficient Huh7 cells (Supplementary Fig. S3h-j). We also evaluated the role of SRCAP, a paralogous protein of EP400, which can also mediate the deposition of H2A.Z on the genome(40,41). The results showed that knocking out the functional subunits SRCAP or ACTR6 of the SRCAP complex in Huh7 cells did not induce the expression of CD47 and PD-L1(Supplementary Fig. S3k-i). These findings demonstrate that BRD8 suppresses CD47 and PD-L1 expression through the EP400 complex, but independent of its acetyltransferase activity and deposition of H2A.Z.

### Targeting BRD8 induces a cell-intrinsic interferon response by upregulating transposable elements expression

To determine how BRD8 regulates CD47 and PD-L1, we first performed RNA-seq profiling of control and BRD8 KO Huh7 cells. Notably, the number of up-regulated genes was far more than that of down-regulated genes in BRD8 depleted cells for the two independent sgRNAs (Supplementary Fig. S4a-c), implying a transcriptional inhibitory function of BRD8. Importantly, Gene ontology (GO) and Kyoto Encyclopedia of Genes and Genomes (KEGG) enrichment analysis of the differentially expressed genes revealed that the common upregulated genes in BRD8-deficient cells were significantly enriched with terms related to IFN response, antigen and immune signaling pathways (Fig. 1h and Supplementary Fig. S3h). Activation of these pathways in BRD8-KO cells was further validated by RT-qPCR experiments, including IFN stimulated genes (ISGs) (e.g., IFIH1), and T cell related-cytokines (e.g., CXCL10) (Fig. 1i). Moreover, major histocompatibility complex class I (MHC I) cell surface expression (e.g., HLA-A, HLA-B and HLA-C) was also elevated in BRD8 deficient cells by RT-qPCR and flow cytometry (Fig. 1i-j). Lastly, western blotting analysis revealed the enhanced TBK1 phosphorylation upon BRD8 loss (Fig. 1k), a marker of IFN pathway activation. Additionally, the expression of tumor immunogenic genes was further upregulated after IFN-γ treatment in BRD8-KO cells according to the RNA-seq data (Supplementary Fig. S4d). These results together demonstrate that BRD8 deficiency triggers cell-intrinsic interferon response and sensitizes cellular immune response to IFN-γ.

We next performed chromatin immunoprecipitation assays followed by DNA sequencing (ChIP-seq) in Huh7 cells stably expressing Flag-tagged BRD8. Notably, among 6,441 high-confidence BRD8 binding peaks, ∼31% located at promoter regions (Supplementary Fig. S4e-f). Interestingly, almost none of the upregulated immune response genes appeared to be direct targets of BRD8, as ChIP-seq experiments with anti-Flag antibody failed to detect BRD8 binding at their promoters (Supplementary Fig. S4g). Moreover, the genes bound by BRD8 at the promoter regions have little correlation with the differentially expressed genes in RNA-seq (Supplementary Fig. S4h-i). These data support that BRD8 indirectly represses these immune response genes.

In a search for the mechanism of BRD8-mediated repression, we conducted ingenuity pathway analysis (IPA) of the RNA-seq data on genes upregulated by BRD8 deletion (Fig. 2a). Poly rI:rC RNA, an artificial double standards RNA (dsRNA) analogue, stood out as one of the top upstream regulators. Considering TIP60 and ING3, the members of BRD8/EP400 complex, their depletion led to de-repression of endogenous retroviral elements in human cancer cells(42,43). We reasoned that BRD8 likely exerts analogous TE regulatory functions in HCC. Strikingly, BRD8 genetic perturbation significantly activated TE transcription in HCC (Fig. 2b). Furthermore, total RNA-seq analysis showed that 283 TEs were significantly up-regulated while only 3 TEs were down-regulated, including long terminal repeat (LTR)-containing endogenous retroviruses (ERVs) and non-LTR elements, such as long interspersed nuclear elements (LINEs) and short interspersed nuclear elements (SINEs) in Huh7 cells upon BRD8 loss (Fig. 2c-d). Accordingly, we detected dsRNA accumulation in BRD8 depleted Huh7 cells with dsRNA specific J2 antibody, but not the case in H2A.Z depleted cells (Fig. 2e-f and Supplementary Fig. S3m). These findings mechanistically link BRD8 deficiency to dsRNA-mediated innate immune activation through de-repression of retrotransposons.

**Fig. 2.**
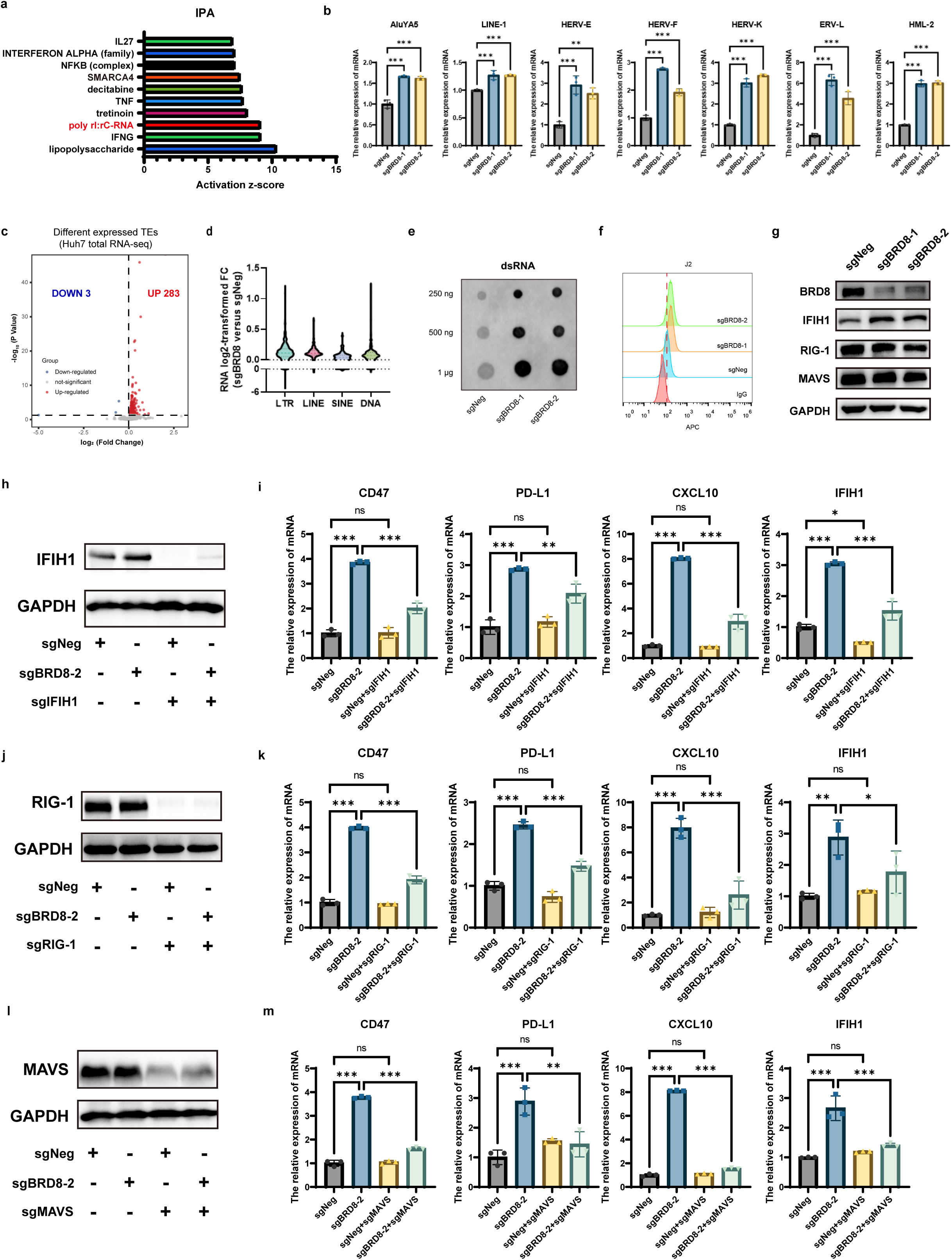
BRD8 negatively regulates transcription of repetitive elements and accumulation of cellular dsRNA. (**a**) Ingenuity pathway analysis (IPA) of upstream regulators of genes activated by BRD8 depletion based on genes in RNA-seq. (**b**) Selected TEs mRNA levels in Huh7 cells upon BRD8 depletion. Results is shown as mean ± SD (n = 3). (**c**) Volcano plot of total RNA-seq analysis in WT and BRD8-deficent Huh7 cells (FDR-adjusted p < 0.05). (**d**) The differentially expressed retroelement classes in (**c**). (**e**) J2-IP dot blot showing dsRNA staining in control and BRD8 knockout Huh7 cells. (**f**) Flow cytometry histograms of dsRNA staining of WT or BRD8 KO Huh7 cells. (**g**) Western Blot of IFIH1, RIG-1 and MAVS in control and BRD8 knockout in Huh7 cells. GAPDH was used as loading control. (**h,j,l**) Western Blot of IFIH1(**h**), RIG-1(**j**), MAVS(**l**) in control, BRD8-KO, DKO in Huh7 cells. GAPDH was used as loading control. (**i,k,m**) RT-qPCR analyses of ISGs in WT, KO and D-KO Huh7 cells. Results are shown as mean ± SD (n=3). ns, not significant, \**p* < 0.05, \*\**p* < 0.01, \*\*\**p* < 0.001, unpaired Student’s *t*-test.

Intracellular dsRNA is recognized by pattern recognition receptors, MDA5 (encoded by IFIH1), RIG-I (encoded by DDX58), and their common downstream MAVS, which are involved in subsequent activation of the IFN pathways(8). Notably, all of these receptors had normal expression levels and IFIH1 was among the upregulated genes in BRD8-deficient cells (Fig. 1i and 2g). To determine whether dsRNA accumulation by BRD8 loss elicits cellular immune responses, we individually disrupted the expression of IFIH1, RIG-1, and MAVS. Importantly, abrogation of the intracellular dsRNA sensors (IFIH1 or RIG-1) significantly diminished the induction of ISGs in BRD8-depleted cells, and depletion of MAVS almost completely counteracted the effect of BRD8 loss, with a level similar to the control (Fig. 2h-m). Since hepatocytes do not express STING(44,45), we do not consider the role of dsRNA-derived dsDNA that connects the cGAS/STING pathway with IFN signaling activation. Collectively, these data demonstrate that BRD8 depletion activates the cell-intrinsic IFN response in liver cancer through the dsRNA-IFIH1/RIG-1-MAVS pathway.

### BRD8 loss redistributes H3K27ac to enable TE activation

Previous studies have shown that components of the BRD8 complex influence TE expression through repressive chromatin marks(42,43). However, global protein levels of the repressive chromatin marks H3K9me3, H4K20me3, and H3K27me3 were unchanged in BRD8-knockout cells (Supplementary Fig. S5a). To determine how BRD8 represses TE expression, we performed assays for transposase-accessible chromatin with sequencing (ATAC-seq) to examine the chromatin landscape. Across the genome, 5,321 sites gained accessibility and 2,536 sites lost accessibility upon BRD8 loss (log2fold change>0.5) (Fig. 3a), consistent with the RNA-seq data showing more upregulated genes than downregulated genes (Supplementary Fig. S4a-b). Importantly, among these sites with increased accessibility, 1,427 were annotated as TEs (ATAC-UP_TE regions) that include SINE, LINE and ERV, while only 463 TE-containing sites showed decreased accessibility (Fig. 3b). Moreover, BRD8-bound regions rarely overlapped with the ATAC-UP_TE regions, and ATAC-seq signal was reduced at BRD8-bound regions, arguing against that BRD8 directly represses the TE expression (Supplementary Fig. S5b-c). These data suggest that BRD8 loss indirectly enhances the chromatin accessibility of TE-containing sites, leading to their de-repression.

**Fig. 3.**
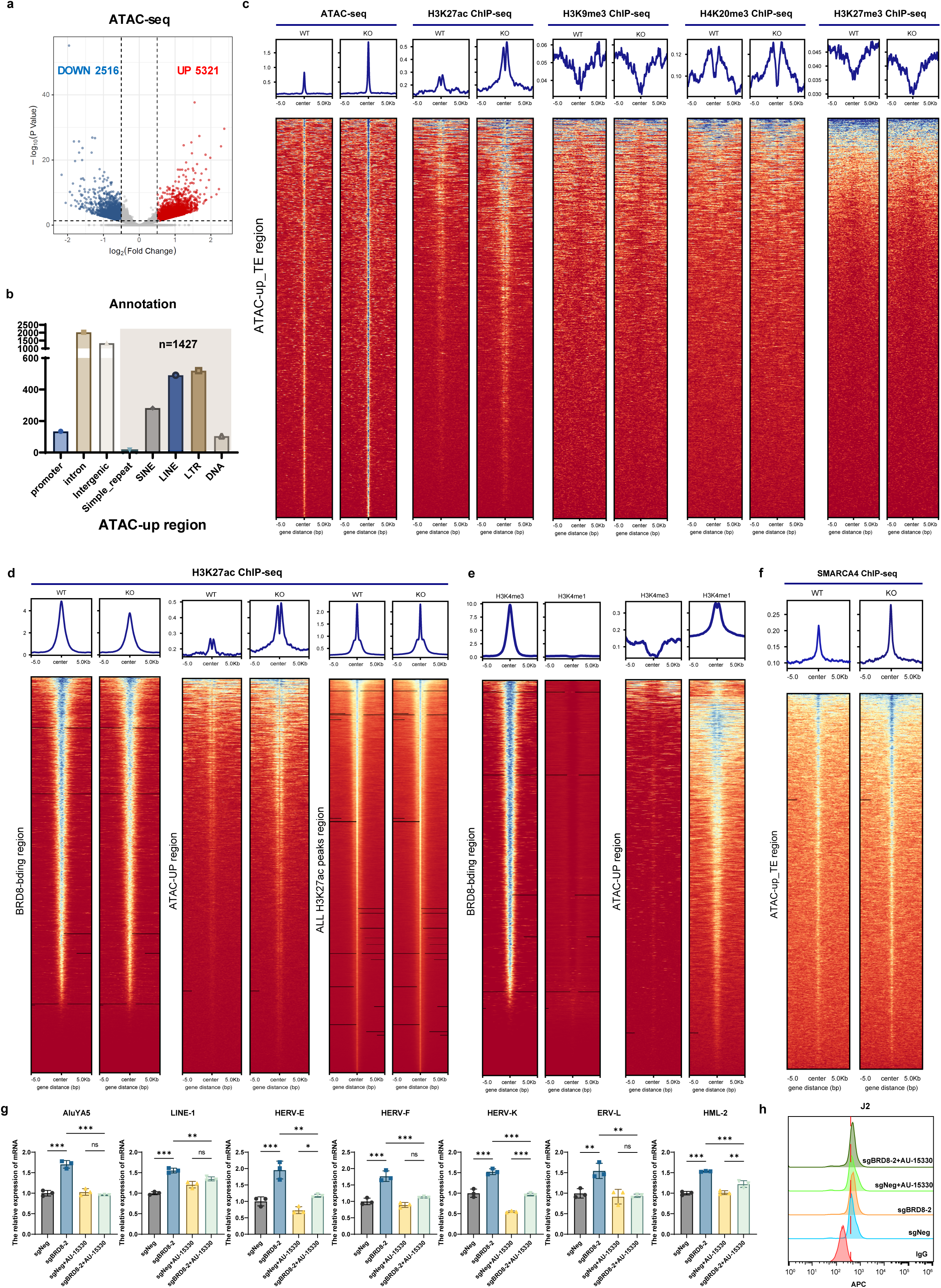
BRD8 regulates accessibility of chromatin and redistribution of H3K27ac occupancy. (**a**) Volcano plot of ATAC-seq analysis. Shown are ATAC-seq all peaks increased (red) and decreased (blue) in BRD8-deficent Huh7 cells (FDR-adjusted p < 0.05). (**b**) The amounts of upregulated repeats in BRD8 knockout Huh7 cells. (**c**) Binding density heatmap of ATAC-seq, H3K27ac, H3K9me3, H4K20me3 and H3K27me3 ChIP-seq signals at ATAC-up_TE region in WT or BRD8-deficent Huh7 cells. (**d**) Binding density heatmap of H3K27ac ChIP-seq signals at BRD8-binding region , ATAC-up region or all H3K27ac peaks region in WT or BRD8-deficent Huh7 cells. (**e**) Binding density heatmap of H3K4me3 or H3K4me1 ChIP-seq signals at BRD8-binding region or ATAC-up region in WT or BRD8-deficent Huh7 cells. (**f**) Binding density heatmap of SMARCA4 ChIP-seq signals at ATAC-up region in WT or BRD8-deficent Huh7 cells. (**g**) RT-qPCR analyses of TEs mRNA levels in Huh7 cells upon BRD8 depletion with or without AU-15330 treatment for 72 h. Results is shown as mean ± SD (n = 3). (**h**) Flow cytometry histograms of dsRNA staining of WT or BRD8 KO Huh7 cells with or without AU-15330 treatment for 72 h. ns, not significant, \**p* < 0.05, \*\**p* < 0.01, \*\*\**p* < 0.001, unpaired Student’s *t*-test.

We further analyzed the changes of histone modifications at the ATAC-UP_TE regions by ChIP-seq. Interestingly, the signals of H3K9me3, H4K20me3, and H3K27me3, repressive marks that are usually considered to be required for TE silencing(12,13,46–49), remained largely unchanged, while the H3K27ac signals, active enhancer marks, robustly increased at these sites in BRD8 deficient cells (Fig. 3c and Supplementary Fig. S5d), linking BRD8 to H3K27ac distribution. Indeed, BRD8 depletion diminished the H3K27ac signals in BRD8-bound regions but minimally affected the global H3K27ac signals, in line with a recent study showing that BRD8 is required for maintenance of histone acetylation(50). In contrast, all the 5,321 sites with increased chromatin accessibility (ATAC-UP regions) covering the 1427 ATAC-UP_TE regions displayed remarkable upregulation of H3K27ac signals (Fig. 3d). Further epigenetic feature analyses revealed that BRD8-bound regions were mainly enriched with H3K4me3 signals while ATAC-UP regions were enriched with H3K4me1 signals (Fig. 3e), representing promoter and poised enhancer regions, respectively. Together, these data indicate that BRD8 loss leads to a redistribution of H3K27ac marks from promoter regions where BRD8 binds to poised enhancer regions, enabling the enhancer activation, thereby de-repressing the TEs within these activated enhancers.

SMARCA4, a core member of the SWI/SNF complex, was also identified as one of the upstream regulators by the IPA of the RNA-seq data (Fig. 2a). Both H3K4me1 and H3K27ac enriched regions were previously reported to enable effective recruitment of SMARAC4 for subsequent transcription activation(51,52). We asked whether SMARCA4 is involved in the TE activation in BRD8 deficient cells. In fact, BRD8 knockout resulted in an increase of SMARCA4 ChIP-seq signals at the 5321 regions with increased chromatin accessibility including the 1427 TE regions (Fig. 3f and Supplementary Fig. S5d). Furthermore, either the SMARCA4 degrader AU-15330 or the enzyme inhibitor BRM014 treatment were able to effectively counteracted the TE activation and upregulation of CD47 and PD-L1 caused by BRD8 loss (Fig. 3g and Supplementary Fig. S5e). In contrast, two other common activator inhibitors, A-485 (EP300/CBP inhibitor) and JQ1 (BRD2/3/4 inhibitor), failed to do so (Supplementary Fig. S5f-g). Taken together, these data demonstrate that BRD8 depletion leads to redistribution of H3K27ac for TE activation, which is in need of SMARCA4.

### Loss of BRD8 induces dsRNA stress that triggers anti-tumor immunity

We next explored biological consequences of BRD8 ablation-induced dsRNA stress, and in particular, whether dsRNA stress-triggered cell-intrinsic immune responses can be harnessed for anti-tumor immunity. To address these questions, we first asked whether our observations made in human HCC cells could be recapitulated in murine HCC cells. Notably, Brd8 depletion resulted in upregulation of retrotransposons, accumulation of dsRNA and subsequent activation of IFN pathways in murine HCC cell line RIL-175 and Hep53.4 (Supplementary Fig. S6), which recapitulates our findings in human HCC cells and reflects the conserved function of BRD8 between humans and mice.

To examine whether Brd8 influences anti-tumor immunity, we inoculated immune-competent C57BL/6J mice subcutaneously and liver orthotopically with RIL175 cells, respectively. Remarkably, Brd8 deficiency significantly suppressed tumor growth, resulting in reduced tumor size and increased animal survival (Fig. 4a-c). Importantly, this effect was not due to impaired cell growth since Brd8 deficiency resulted in no compromised growth of RIL-175 cells *in vitro,* and had minimal effect on HCC tumor growth and animal survival in immunodeficient NOD.Cg-Prkdc^scid^ Il2rg^tm1Wjl^/SzJ (NSG) mouse model (Supplementary Fig. S7a-c).

**Fig. 4.**
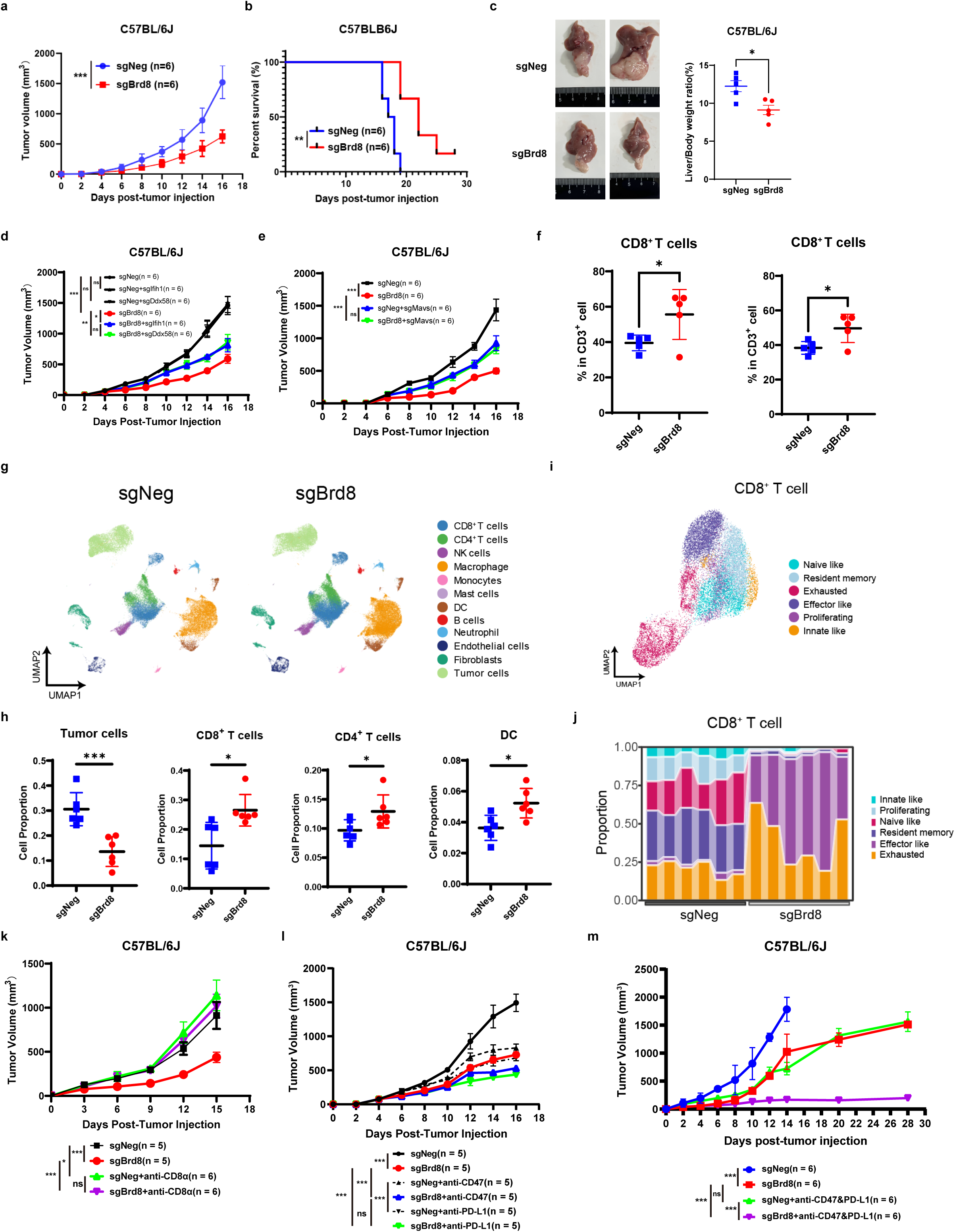
Depletion of Brd8 induces anti-tumor immunity and sensitizes tumors to ICB therapy. (**a**) Tumor growth curves of C57BL/6J mice subcutaneously injected with 2,000,000 control (n = 6) or Brd8 sgRNA (n = 6) RIL-175 cells. (**b**) Survival curves of C57BL/6J mice injected with control (n = 6) or Brd8 sgRNA (n = 6) RIL-175 cells. (**c**) Representative image and liver/body weight ratio of HCC orthotopic C57BL/6J mice model with control (n = 5) or Brd8 sgRNA (n = 5) RIL-175 cells. (**d** and **e**) Tumor growth curves of C57BL/6J mice subcutaneously injected with RIL-175 cells of the indicated genotypes. d, control only (n = 6), Brd8 sgRNA only (n = 6), control and Ifih1 sgRNA (n = 6), Brd8 sgRNA and Ifih1 sgRNA (n = 6), control and Ddx58 sgRNA (n = 6), Brd8 sgRNA and Ddx58 sgRNA (n = 6); e, control only (n = 6), Brd8 sgRNA only (n = 6), control and Mavs sgRNA (n = 6), Brd8 sgRNA and Mavs sgRNA (n = 6). (**f**) Percentage of tumor-infiltrating lymphocytes in C57BL/6 mice injected with control or Brd8 sgRNA RIL-175 cells described in (a, left) and (c, right). Each dot represents the infiltrating lymphocytes in a single tumor. (**g**) Uniform Manifold Approximation and Projection (UMAP) visualization of cells from control and sgBrd8 tumors and subjected to scRNA sequencing. (**h**) Quantitation of the proportion of various types of cells in control and sgBrd8 tumors. (**i**) UMAP of CD8^+^ T cells from control and sgBrd8 tumors and subjected to scRNA sequencing. (**j**) bar plot quantification of types of CD8^+^ T cells in control and sgBrd8 tumors. (**k**) Tumor growth curves of in C57BL/6 mice treated with isotype control or anti-mouse CD8α antibody before the injection of the RIL-175 Brd8-KO cells described in (a). (**l**) Tumor growth curves of in C57BL/6 mice subcutaneously injected with RIL-175 Brd8-KO cells reconstituted as in (a) and subsequently treated with isotype control, anti-mouse CD47 antibody or anti-mouse PD-L1 antibody. (**m**) Tumor growth curves of in C57BL/6 mice subcutaneously injected with RIL-175 Brd8-KO cells reconstituted as in (a) and subsequently treated with isotype control or anti-mouse CD47 plus PD-L1 bis-antibodies. ns, not significant, \**p* < 0.05, \*\**p* < 0.01, \*\*\**p* < 0.001, unpaired Student’s *t*-test.

To further determine whether the tumor growth phenotype is due to BRD8 deficiency-induced dsRNA stress, we individually deleted dsRNA sensors Ifih1 or Ddx58 and their common downstream Mavs in Brd8-KO RIL-175 cells. As expected, depletion of either Ifih1 or Ddx58 partially rescued the tumor-forming ability (Fig. 4d). Moreover, abrogation of Mavs fully rescued the *in vivo* tumor growth phenotype of Brd8 deficient cells (Fig. 4e). These data demonstrate that Brd8 deficiency elicits a dsRNA-dependent anti-tumor immune response.

### Brd8 abrogation remodels TME and sensitizes tumors to ICB therapy

To further elucidate the mechanism connecting Brd8 inhibition with enhanced anti-tumor immunity, we investigated the effects of Brd8 depletion on the tumor microenvironment (TME). Flow cytometry analysis revealed a significant increase of CD8+ cytotoxic T cells of intra-tumoral immune cells in both subcutaneous and liver orthotopic tumor models (Fig. 4f and Supplementary Fig. S7d-f). Considering individual variation in the orthotopic model, we next performed single-cell RNA sequencing (scRNA-seq) on subcutaneous tumors. As expected, Brd8-KO tumors contained significant decreased tumor cells, but increased CD8+ cytotoxic T cells, CD4+ help T cells and dendritic cells (Fig. 4g-h). Moreover, tumor-infiltrating CD8+ T cells clustered into six clusters based on well-established marker genes: naive-like, resident memory, effector-like, exhausted, innate-like, and proliferating. We found that the majority of CD8+ T cells from control tumors clustered in the innate-like cluster, naive-like cluster and resident memory cluster, while CD8+ T cells from Brd8-deficiency tumors showed tremendously increased density in the effector-like cluster and exhausted cluster (Fig. 4i-j). This altered cellular states of CD8+ T cells revealed insights into CD8+ T cell differentiation states in our model. Notably, the scRNA-seq data confirmed our flow cytometry data and indicated that CD8+ T cells are essential effectors for the observed anti-tumor immune response. Indeed, depletion of CD8+ T cells fully rescued the growth of Brd8-deficient tumors (Fig. 4k and Supplementary Fig. S7g). These findings demonstrate that Brd8 loss enhances tumor immunogenicity and cytotoxic T cell-mediated tumor rejection through remodeling the TME.

PD-L1 expression is significantly associated with clinical response to anti-PD-1/PD-L1 antibodies(6,7). The fact that PD-L1 expression was upregulated in Brd8-KO cells and ratio of exhausted T cells was increased in Brd8-deficiency tumors prompted us to assess whether PD-L1 blockade would synergize with Brd8 ablation to further enhance anti-tumor immunity. Indeed, Brd8 depletion dramatically enhanced the sensitivity of RIL-175 tumors to anti–PD-L1 therapy. As another important immune checkpoint(53,54), we speculated that upregulated CD47 in Brd8-KO cells should also has the similar effect on anti–CD47 therapy. Although the effect was not as well as anti–PD-L1 therapy when used alone, the sensitivity of RIL-175 tumors to anti–CD47 therapy was significantly improved upon Brd8 loss (Fig. 4l and Supplementary Fig. S7h). Previous research has revealed bispecific antibody of CD47/PD-L1 could maximizes antitumor immunity(55). As a proof of concept, we assessed whether dual anti-CD47/PD-L1 antibody could further enhance anti-tumor immunity compared to anti-CD47 or anti-PD-L1 therapy alone in Brd8 deficient tumors. Remarkably, treatment of Brd8-deficient RIL-175 tumors with dual anti-CD47/PD-L1 antibody almost completely abrogated the tumor forming ability (Fig. 4m and Supplementary Fig. S7i). Taken together, these results demonstrate that Brd8 deficiency remodels the TME to enhance tumor immunogenicity and synergizes ICB therapy to abrogate tumor growth.

### Targeting bromodomain of BRD8 induces cell-intrinsic immunity and anti-tumor effect

We identified the bromodomain of BRD8 as the most selective CD47/PD-L1 regulator by domain-focused CRISPR screening for 52 bromodomains within 39 human proteins (Fig. 5a). Previous studies showed that the BD domain of BRD8 has an independent evolutionary branch(25,27). To interrogate the role of the bromodomain in BRD8, we performed rescue experiments with WT or bromodomain deletion form of BRD8(Supplementary Fig. S8a-b). Notably, CRSIPR-editing resistant full-length BRD8 completely restored the expression levels of TE and ISGs while bromodomain-deleted BRD8 failed in BRD8 deficient cells (Fig. 5b-c and Supplementary Fig. S8c). These results demonstrate that the bromodomain is crucial for the function of BRD8.

**Fig. 5.**
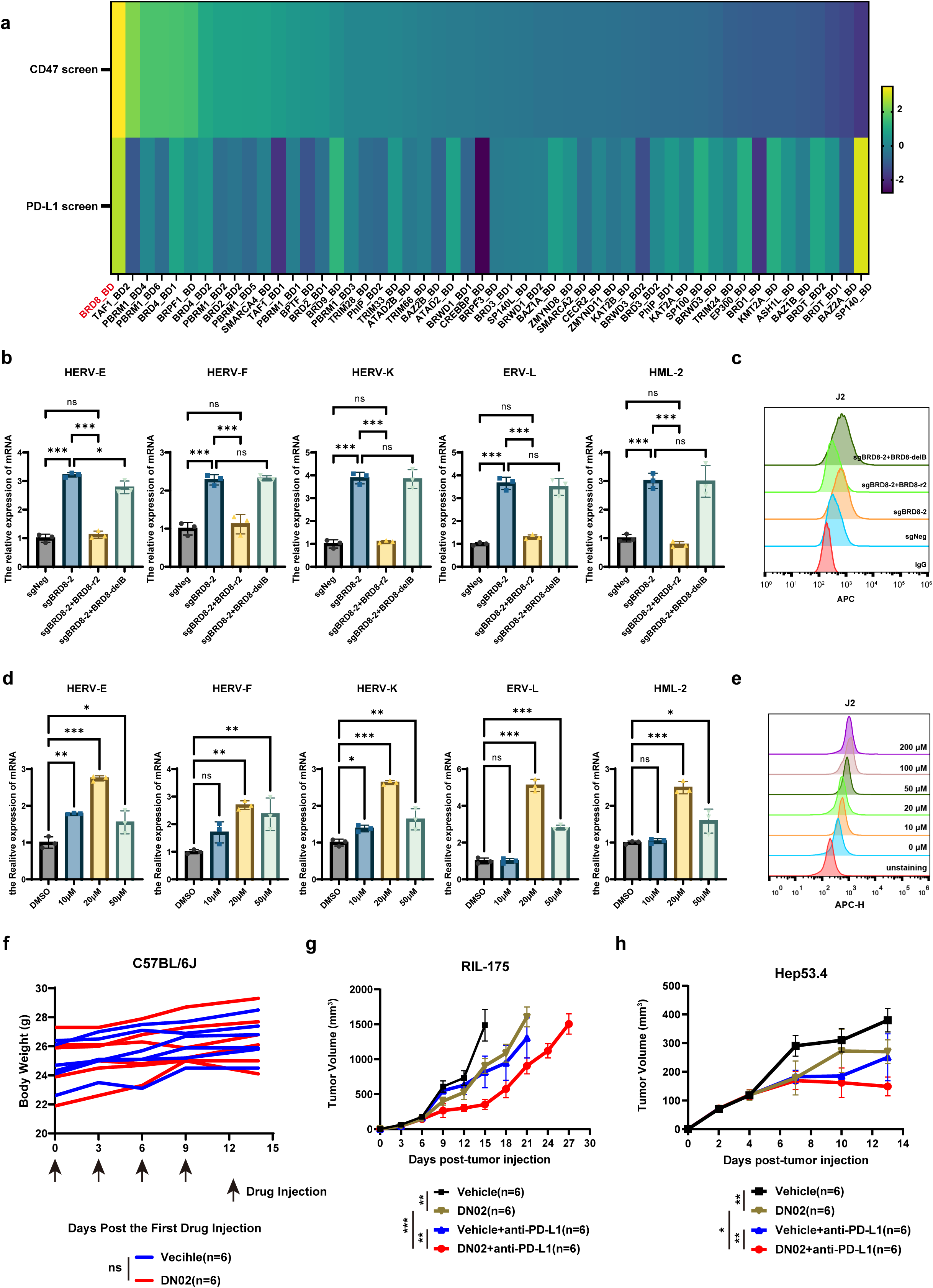
The bromodomain of BRD8 is selectively essential for its function. (**a**) Heat map showing CRISPR screen results for the bromodomain-containing proteins. (**b**) RT-qPCR analyses of TEs mRNA levels in Huh7 BRD8-KO cells reconstituted with CRISPR-resistance (BRD8-r2) or BD deletion (BRD8-delB) BRD8 cDNA. Results is shown as mean ± SD (n = 3). (**c**) Flow cytometry histograms of dsRNA staining of Huh7 cells described in (b). (**d**) RT-qPCR analyses of TEs mRNA levels in Huh7 cells with different dosages of DN02 treatment for 72 h. Results is shown as mean ± SD (n = 3). (**e**) Flow cytometry histograms of dsRNA staining of Huh7 cells described in (d). (**f**) Body weights of C57BL/6 mice subcutaneously injected with RIL-175 cells subsequently treated with vehicle or DN02 (40 mg/kg; every three days; four times). (**g**) Tumor growth curves of in C57BL/6 mice described in (f) and subsequently treated with isotype control or anti-mouse PD-L1 antibody. (**h**) Tumor growth curves of in C57BL/6 mice subcutaneously injected with 5,000,000 Hep53.4 cells subsequently treated with vehicle or DN02 plus isotype control or anti-mouse PD-L1 antibody. ns, not significant, \**p* < 0.05, \*\**p* < 0.01, \*\*\**p* < 0.001, unpaired Student’s *t*-test.

DN02 represents the first reported selective chemical probe targeting the BRD8 bromodomain(56). Despite its initial characterization by Remillard et al, no functional applications have been reported. To establish foundational pharmacodynamic parameters, we first determined cytotoxic potential of DN02 in HCC cells, revealing IC50 values of 74.19 μM (Huh7) and 193.1 μM (RIL-175) through dose-response profiling (Supplementary Fig. S8d). Importantly, DN02 treatments led to induction of TEs, accumulation of cytoplasmic dsRNA and subsequent upregulation of ISGs in Huh7 cells (Fig. 5d-e and Supplementary Fig. S8f-g), phenocopying the BRD8 knockout effects. Of note, DN02 treatments exhibited no effect on BRD8 protein abundance (Supplementary Fig. S8e). We next evaluated translational potential of DN02 through comprehensive in vivo studies. Initial safety assessments in C57BL/6J mice receiving intraperitoneal injections showed preserved locomotor activity and stable body mass without mortality, confirming acceptable therapeutic tolerability (Fig. 5f). Anti-tumor efficacy testing revealed immuno-dependent activity: DN02 monotherapy significantly inhibited tumor progression in C57BL/6J mice, but it failed to suppress RIL-175 tumor growth in NSG mice (Supplementary Fig. S8h-i). We further tested combination therapy in two HCC models (RIL-175 and Hep53.4). Notably, co-administration of DN02 with anti-PD-L1 antibody significantly enhanced tumor suppression compared to monotherapies (Fig. 5g-h). Thus, these results suggest the potential therapeutic value of DN02 for HCC.

### BRD8 expression is negatively correlated with IFN signatures, immune infiltration and associated with poor clinical outcome in human cancer patient datasets

To determine the clinical and translational significance of our findings in human cancer patients, we first performed immunohistochemistry experiments on two HCC tumor microarrays (TMAs) from Zhongshan Hospital. Notably, the HCC patients expressing high levels of BRD8 protein tended to have low PD-L1 protein expression in tumor cells and a reduced number of CD8+ T cells in tumors, displaying a negative correlation, respectively (Fig. 6a-c). Moreover, we examined the expression of BRD8 in liver cancer from The Cancer Genome Atlas (TCGA) dataset. There is a significant different expression of BRD8 between the tumor and normal samples in TCGA-LIHC (Supplementary Fig. S9a). And BRD8 expression is negatively correlated with interferon and other immune-related signaling pathways (Fig. 6d). Further single sample gene set enrichment analysis(ssGSEA) showed that in liver cancer samples with low BRD8 expression, the infiltrations of activated CD8+ T cells and memory CD8+ T cells, were significantly higher compared to those with high BRD8 expression (Fig. 6e). Interestingly, this correlation was also observed in several other tumor types (Supplementary Fig. S9b-f). Additionally, analysis of RNA-seq data for TILs in patients with liver cancer(57) or non-small cell lung cancer (NSCLC)(58) treated with an anti–PD-1 antibody revealed that BRD8 levels were significantly higher in non-responders compared with responders (Fig. 6f). These results collectively suggest that cancer patients with low BRD8 expression tend to have activated interferon signals and immune microenvironment, and can benefit from immunotherapy.

**Fig. 6.**
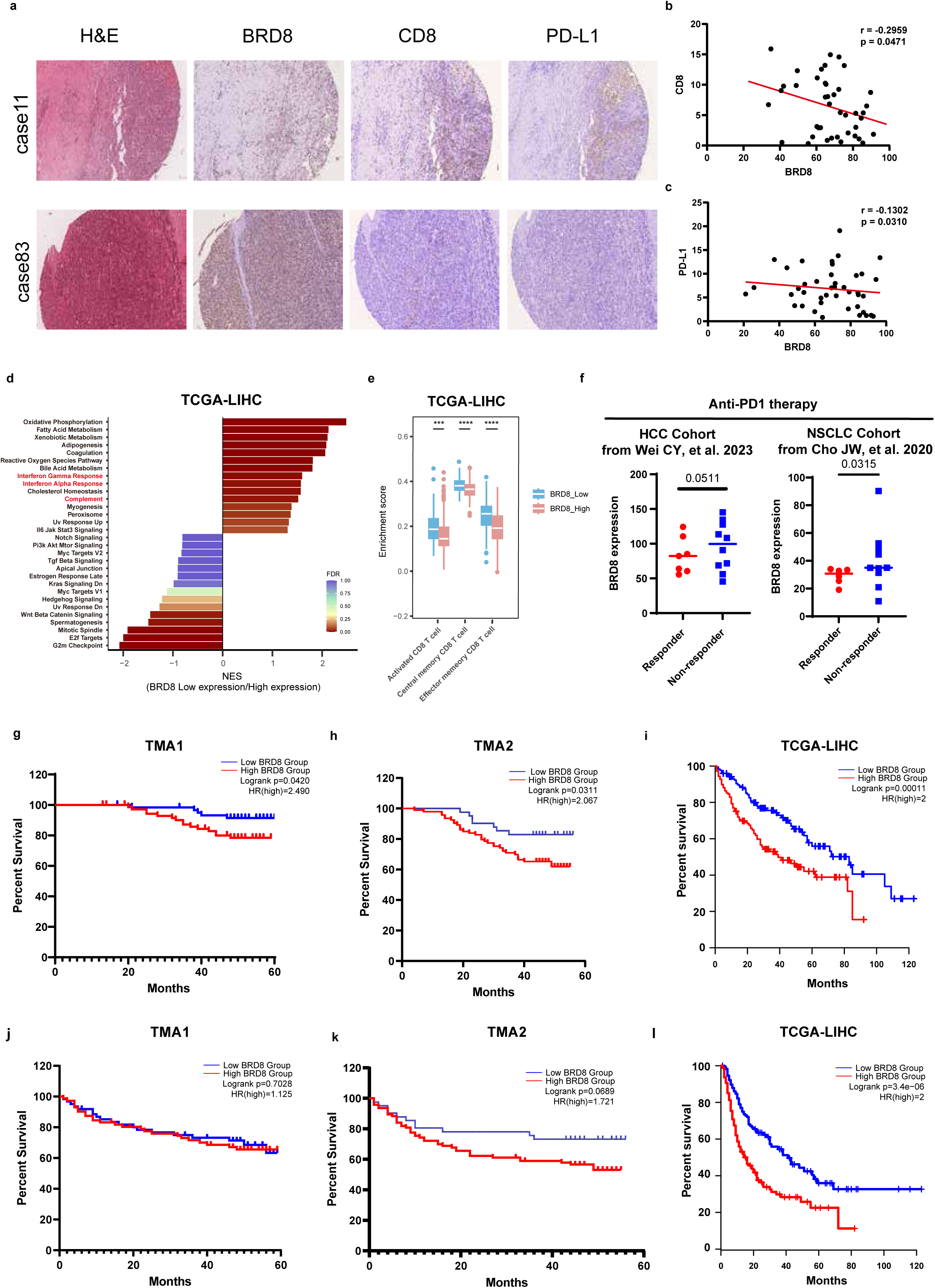
Analysis of the clinical outcome of cancer patients with BRD8 expression. (**a**) Two representative HCC patients (Case 11 and Case 83) showing different levels of BRD8, CD8 and PD-L1 expression (100×). Patient 1 shows BRD8^low^CD8^high^PD-L1^high^ and Patient 2 shows BRD8^high^CD8^low^PD-L1^low^. (**b,c**) The correlations between BRD8 expression and CD8(b) or PD-L1(c) expression were calculated on protein level in HCC cohort. (**d,e**) GSEA(d) and ssGSEA(e) based on RNA-seq results of BRD8 high and low expression patients in TCGA-LIHC dataset. (**f**) BRD8 levels prior to treatment with anti-PD-1 checkpoint inhibitors in non-responder and responder groups of patients from the referenced cohorts. p values were calculated by unequal variance t test. (**g-l**) The overall survival(g-i) or disease-free survival(j-l) curves of BRD8-low and BRD8-high in the TMA1(g,j), TMA2(h,k) and TCGA-LIHC(i,l) dataset.

We then examined whether BRD8 expression level in tumors correlates with clinical outcome. Indeed, HCC patients with high BRD8 levels in tumor samples experienced significantly shorter overall survival (OS) than those with low BRD8 expression in our two independent TMA cohorts (Fig. 6g-h). Low BRD8 expression correlated with low Hazard Ratio (HR) of disease-free survival, despite there is no significance (Fig. 6j-k). Moreover, according to the TCGA datasets, the overall survival and disease-free survival of patients with high BRD8 expression were significantly lower than those with low BRD8 expression, consistent with our datasets. (Fig. 6i, l and Supplementary Fig. S8g). Taken together, these findings suggest that BRD8 may become an effective immuno-oncology treatment target and a biomarker to predict immunotherapy responsiveness and patient prognosis in human cancers.

## Discussion

Through domain-focused CRISPR screening, we identify BRD8 as a top negative regulator of the immune checkpoint CD47 and PD-L1, operating via the BRD8/EP400 chromatin remodeling complex independent of its acetyltransferase activity and deposition of H2A.Z. Further mechanistic study reveals that BRD8 ablation redistributes H3K27ac to repetitive genomic elements, derepressing endogenous retrotransposons (LTRs, LINEs, SINEs) and triggering dsRNA accumulation. This dsRNA stress activates the MDA5/RIG-I-MAVS axis, culminating in type I IFN signaling induction, MHC-I upregulation, and CD8+ T cell-mediated tumor rejection. Moreover, BRD8 loss sensitizes tumors to ICB therapies by remodeling the tumor microenvironment (TME) toward cytotoxic T cell infiltration. The first-in-class BRD8 bromodomain inhibitor DN02 treatment elicits immunomodulatory and anti-tumor effects, mimicking BRD8 loss. Therefore, this study establishes BRD8/EP400 complex as a master epigenetic suppressor of tumor-intrinsic immunogenicity in HCC and proposes the translational potential of the BRD8 bromodomain-targeting inhibitor as a strategy to enhance response to immunotherapy for immune-cold liver cancer patients.

BRD8 has been reported to function as an oncogene in HCC(24), GBM(32), and colon cancer(23,25). In these models, BRD8 influences tumor progression mainly by regulating cell division. Moreover, BRD8 was also reported to be associated with immunity in HCC(28), NSCLC(29), renal cell cancer(59) and colon cancer(26) with little mechanistic insights. In our models, BRD8 depletion displayed minimal effect on cell proliferation in vitro and in vivo. Importantly, our study reveals the detailed mechanism through which BRD8 engages in the regulation of tumor intrinsic immunogenicity. BRD8 thus joins EZH2 and LSD1 as a chromatin-based tumor-intrinsic immune regulator, but links H3K27ac occupancy dysregulation to TE-driven immunogenicity, rather than affecting TE-associated repressive marks (H3K9me3, H4K20me3, H3K27me3). In line with our study, it has been reported that increased enrichment of H3K27ac enables efficient TEs activation in mouse embryonic stem cells(60) and GBM(61).

BRD8 depletion induces a redistribution of H3K27ac from promoter regions to poised enhancer regions containing TEs, revealing a novel mechanism for TE activation. This aligns with the previously established paradigm linking histone modification redistribution to TE control, exemplified by Menin perturbation driving MLL1-mediated H3K4me3 relocation to TEs(62). The spatial reorganization of specific histone modifications across the different genomic regions represents a fundamental epigenetic mechanism for dynamic gene regulation, as demonstrated by H3K4me3 redistribution during sperm maturation(63) and H3K9ac relocation during yeast starvation(64). This process—whereby a mark is erased from one genomic compartment and deposited at another—enables rapid, locus-specific functional reprogramming without requiring de novo protein synthesis. Such redistribution appears critical for epigenetic regulation across diverse physiological and pathological contexts. BRD8 is thought to exerts its function through directly binding to acetylated histones with its bromodomain(25,27). Indeed, BRD8 depletion reduces H3K27ac signals at its binding regions but minimally affects global H3K27ac enrichment, consistent with prior observations in mouse embryonic stem cells(50). Moreover, unlike WT BRD8, the bromodomain-deleted form fails to rescue the IFN-induced phenotype, highlighting its crucial role. More importantly, treatment of DN02, a first-in-class selective BRD8 bromodomain probe, mimicks the effect of BRD8 genetic ablation by inducing dsRNA stress and IFN responses, although to a lesser extent. To our knowledge, we test the safety and effective anti-tumor function of DN02 in murine tumor models for the first time, revealing the translational potential of BRD8 bromodomain inhibitors. However, the overall efficacy of DN02 is limited compared with knockout of BRD8, and thus safer and more effective BRD8 bromodomain inhibitors warrant to be developed in the future.

TE-derived dsRNA can activate cell-intrinsic IFN signaling pathway known as “viral mimicry response” that promotes cell death(8). Meanwhile, this phenomenon also induces the expression of immune checkpoints such as PD-L1 and CD47 which would exhaust cytotoxic T cells and macrophage, respectively. Indeed, our scRNA-seq data reveals the increase of exhausted T cells in BRD8 deficient tumors. As a proof of concept, we assess the anti-tumor effect of BRD8-loss induced viral mimicry combined with ICB therapies. Strikingly, BRD8 depletion sensitizes murine tumors to anti-PD-L1 therapy and anti-CD47 therapy. Particularly, dual anti-PD-L1/CD47 therapy almost completely abolishes the tumor-forming ability of BRD8 deficient murine HCC cells, maximizing the anti-tumor immunity compared with anti-PD-L1 or anti-CD47 alone. Thus, these findings provide a rationale of combining dual anti-PD-L1/CD47 therapy to optimize tumor immunotherapy.

In summary, our study presents BRD8/EP400 complex as a major epigenetic repressor of tumor-intrinsic immunogenicity and provides a strong rationale for targeting BRD8 bromodomain alone or in combination with immunotherapy for liver cancer patients.

## Methods

### Cell culture

Human HCC cell lines Huh7, HepG2, MHCC-97L cells were obtained from Liver Cancer Institute, Fudan University (Shanghai, China). Human embryonic kidney cell line 293T (HEK293T) was obtained from ATCC. Murine HCC cell lines Hep53.4 was purchased from Meisen CTCC (Shanghai, China), and RIL-175 was a gift from Jun Yu (The Chinese University of Hong Kong, China). These cells were maintained in DMEM media (Gibco) or RPMI-1640 media (Gibco) with 10% fetal bovine serum and 1% penicillin-streptomycin solution. All cell lines were authenticated by short tandem repeat (STR) profiling and periodically tested and shown to be mycoplasma negative. Cells were cultured at 37°C with 5% (v/v) of CO_2_.

### Clinical Tissue Samples

Two independent HCC tissue microarrays (TMAs) were utilized in this study. All specimens were obtained from HCC patients who were scheduled to undergo surgery at Zhongshan Hospital, Fudan University from July 2016 to December 2017. A total of 134 formalin-fixed paraffin embedded HCC tissues (containing tumor and paratumor compartments) were included in TMA1, and 135 were included in TMA2. All human samples were anonymously coded in accordance with local ethical guidelines (as stipulated by the Declaration of Helsinki). Written informed consent was obtained from each patient, and the study protocol was approved by the Research Ethics Committee of Zhongshan Hospital, Fudan University (Y2022-473).

### Animal Study

The wild-type C57BL/6J mice (6–8 weeks, male) were purchased from the GemPharmatech (Nanjing, China) and NOD-SCID Il2rγ^−/−^ (NSG) mice (6–8 weeks, male) were obtained from Shanghai Model Organisms Center, Inc. (Shanghai, China). All Animals were housed in specific-pathogen-free facilities at the Department of Laboratory Animal Science of Fudan University and all animal experiment protocols were approved by the Institutional Animal Care and Use Committee of Fudan University (DSF-2024-057).

### Plasmid construction

All Cas9-expressing cell lines used in this study were constructed through lentiviral transduction with a 5′-Flag-tagged human-codon optimized Cas9 by plasmid LentiV_Cas9_puro (Addgene, 108100). Individual sgRNAs were cloned into lentiviral expression vectors with an optimized sgRNA scaffold backbone plasmid LRG2.1 (Addgene, 108098) or LRB2.1. The LRB2.1 vector was derived from LRG2.1 by replacing the original GFP with BFP using the In-Fusion cloning method. Briefly, for each individual sgRNA construct, two reverse complimentary sgRNA oligonucleotides with added 5’ overhangs of CACC or AAAC respectively, were annealed and then ligation into the BsmbI-digested LRG2.1 by using T4 DNA ligase. The vector LRB2.1 and LRG2.1 were used infection two sgRNAs together in the same cell. Sequences of all sgRNAs and used for this study are provided in Supplementary Table 6. The vector LentiV_Blast was derived from LentiV_Cas9_Blast (Addgene, 125592) removal of Cas9 by using restriction endonuclease, and used to express BRD8 cDNAs for rescue experiments. Full length wild-type *BRD8* cDNA was amplified from Huh7 cell line by PCR. A HA or FLAG tag was added into *C* terminus of *BRD8* cDNA. CRISPR-resistant synonymous mutant and bromodomain-deletion *BRD8* cDNAs as indicated were amplified from wild-type cDNA and cloned into the LentiV_Blast vector using the In-Fusion cloning system. All of constructs were transformation by using the Stbl3 bacteria strain.

### Lentivirus transduction and infection

Lentivirus was produced in HEK293T cells by transfecting lentiviral plasmids with packaging helper plasmid (psPAX2, Addgene 12260) and envelope helper plasmid (pMD2.G Addgene 12259) using polyethylenimine linear (PEI 40000) transfection reagent. In brief, HEK293T cells were seeded in 10 cm dishes. On the day of cell are ∼90-100% confluent, 10 μg of lentiviral plasmid DNA, 7.5 μg of psPAX2, 5 μg of pMD2.G and 80 μl of 1 mg ml^-1^ PEI were mixed and incubated for 10-15 mins in OPTI-MEM medium for each plate, and then added mixture into HEK293T cells. After 6-8h both transfections, a fresh medium change was performed and the virus-containing supernatant was collected and pooled at 24h and 48h after transfection. For lentivirus infection, virus plus 8 mg ml^-1^ polybrene were added into target cells with ∼40-50% confluent and incubate for 24h, and then medium was changed. At 48h after infection, cells were detected GFP or selected by 2 μg ml^-1^ puromycin (for 5 days) or 10 μg ml^-1^ blasticidin (for 10 days).

### Pooled CRISPR sgRNA library screen and analysis

The domain-focused sgRNA library targeting human chromatin remodeling factors and the screening protocol were performed as described previously. Briefly, Cas9 was stably expressed in Huh7 cells through the transduction of the LentiV_Cas9_puro vector. Lentivirus of the pooled sgRNA library was produced as described above. To ensure transduction of a single sgRNA per cell, Huh7-Cas9 cells were transduced with the pooled sgRNA library.at a low multiplicity of infection (MOI < 0.3). ∼5 million cells were infected in total to yield 1000 x coverage of the sgRNA library in the GFP+ population to maintain the representation of sgRNAs. The GFP-positive cells were sorted by flow cytometry on day 2 post-infection. Transduced cells were cultured for an additional 10 days (total 12 days post-infection). After that, these cells were divided into two groups with or without IFNγ treatment for 48 h. On day 14 post-infection, two groups of cells were further divided into four groups and were stained with APC-CD47 or APC-PD-L1, respectively. Cells sorted into CD47/PD-L1 high and CD47/PD-L1 low populations by FACS. Genomic DNA was extracted from these samples by phenol/chloroform extractions per standard methods. Then, gDNAs were amplified with Phanta Flash Master Mix Polymerase (Vazyme P520). PCR reactions were then pooled for each sample and column purified with QIAGEN PCR purification kit (QIAGEN 28106). Purified reactions were used to Illumina MiSeq library construction and sequencing. Pair-end reads were trimmed at using Cutadapt (v3.5) to remove adaptor. After trimming, the number of reads per sgRNA was acquired using MAGeCK (v0.5.9) -count function. Read counts were normalized using the total read counts. For the comparison analysis of the sorted populations (CD47 high and CD47 low, PD-L1 high and low, with or without IFN-γ), MAGeCK -test function was used and FDR < 0.05 was considered to be statistically significant. Normalized read counts of sgRNAs were log_2_ transformed and plot in RStudio software. The normalized sgRNA read counts data of screening are provided in Supplementary Table 1.

### Quantification PCR with reverse transcription

Total RNA for each sample was extracted using TRIzoI (Thermo Fisher Scientific). Briefly, cells were lysed in 1 ml TRlzol for 5 min at room temperature, followed by adding 200 μl chloroform and incubating for 5 min at room temperature. Upper aqueous phase was separated by centrifuging at 13,000 r.p.m. for 15 min at 4°C. An equal volume of isopropanol was adding and pellet was precipitated by centrifuging at 13,000 r.p.m. for 10 min at 4°C. RNA was washed with 75% ethanol twice and then dissolved in RNase-free water. The extracted RNA was reversely transcribed into cDNA using the PrimeScript RT Reagent Kit (Takara RR037B) according to the manufacturer’s instructions, with the following modifications: 1 μg of RNA samples were first denatured at 65°C for 5 min and cooled down on ice for 2 min before the addition of buffer, primers and reverse transcriptase. The reverse transcription reaction is as follows: 25 ° C for 5 min, 37 ° C for 45 min, 85 ° C for 5 s. The obtained cDNA samples were diluted and used for real-time quantitative PCR (RT-qPCR). SYBR green (Roche, cat#04887352001) and gene specific primers with sequences listed in Table S2 were used for PCR amplification and detection on an ABI QuantStudio 6 Flex system (Thremofisher). All the plotted gene expression data are normalized to *GAPDH*.

### Western blotting

Cells in culture were washed with ice-cold PBS twice to completely remove residual medium and resuspended in RIPA cell lysis buffer with protease inhibitor (Roche, 04693132001) and phosphatase inhibitor (Roche, 04906837001) on ice. Lysate was incubated for 30 min on ice and centrifuged at 14,000g for 15 min. The supernatants were collected, quantified using BCA assay and then mixed with 5× SDS loading buffer and boiled for 10 min at 95 °C. Equal volume and equal quantity of protein samples were subjected to SDS-PAGE and transferred to a nitrocellulose membrane (Bio-Rad, 162-0097). Membranes were blocked with 5% not-fat milk in TBST for 1h and incubated with appropriate primary antibodies at 4°C overnight. On next day, membranes were washed with TBST for 10 min three times and incubated HRP-conjugated secondary antibodies at room temperature for 1h. The membranes were washed again and ECL was used for detection by ChemiDoc MP (Bio-Rad).

### Flow Cytometry

For detecting proteins on the cell membrane, collected cells were washed with PBS and stained with anti-CD47 (BioLegend, 323123), anti–PD-L1 (BioLegend, 329708), and anti–HLA-A, B, C (BioLegend, 311410) antibodies diluted with PBS. The fluorescence levels were compared with isotype control antibodies. For intracellular flow cytometry, collected cells were fixed with 4% formaldehyde in PBS and permeabilized with 0.1% Triton X-100 in PBS. The cells were then incubated with primary antibody against dsRNA (J2; SCICONS, no. 10010500) or isotype controls antibody for 1 h. After that, the secondary goat anti-mouse IgG (H+L) conjugated with Alexa Fluor 647 (Thermo Fisher Scientific, A32728) was added to the cells and incubated for 30 mins.

Single-cell suspensions of tumors were prepared using a gentleMACS tissue dissociation system. The cells were incubated with primary antibody against BV510-mCD45 (BioLegend, 103137), PE-cy7-mCD3 (Thermo Fisher Scientific, 25-0031-82), PerCP-Cy5.5-mCD8 (BioLegend, 100734), FITC-mCD4 (Thermo Fisher Scientific, 11-0042-85), BV421-mCD25 (BioLegend, 102043), PE-mFoxp3 (Thermo Fisher Scientific, 12-5773-82), BV786-mCD11b (Thermo Fisher Scientific, 417-0112-82), BUV495-mF4/80 (BD Biosciences, 565614), APC-mNK1.1 (BioLegend, 108710) for staining. All stained cells were analyzed on an Attune™ NxT Flow Cytometer. Data were analyzed using the FlowJo software.

### Animal experiments

For subcutaneous model, about 2 ×10^6^ WT and Brd8-KO RIL-175 or 5 ×10^6^ WT and Brd8-KO Hep53.4 cells were resuspended in PBS and injected subcutaneously into the right flank of recipient C57BL/6 mice. For BRD8 inhibitor treatment, C57BL/6 and NSG mice were treated with 40 mg/kg DN02 twice per week for a total of 2 weeks after tumor engraftment on day 4. For immunotherapy treatment, C57BL/6 mice were treated with 200 µg anti-PD-L1, 200 µg anti-CD47, 100 µg anti-CD47+PD-L1 antibodies or isotype control twice a week on day 7 after tumor injection and continuing four times. For CD8+ cells depletion, mice were given 200µg of anti-CD8 or rat isotype control via IP starting on day -3 relative to tumor injection, and then every m3 days until completion of the experiment. All tumor volumes were measured by digital calipers and calculated using the formula: (width * width * length)/2. Mice were sacrificed when tumors reached 1500 mm^3^ or upon ulceration/bleeding. Tumor growth and survival data were plotted in GraphPad Prism (v9.5).

For orthotopic model, about 5 ×10^5^ WT and Brd8-KO RIL-175 cells were resuspended in 50 μl serum-free DMEM medium and injected into the left lobe of the liver of anesthetized 6-week-old male C57BL/6 J mice. The mice were sacrificed 21 days after the surgery.

### IP-MS/IP-WB

About 100 × 10^6^ HuH-7 cells were washed with ice cold PBS twice, lysed with lysis buffer A (10 mM HEPES pH 7.5, 1.5 mM MgCl2, 10 mM KCl, 1 mM DTT, and 1 mM PMSF) for 10 mins on ice and then resuspended with nuclei lysis buffer C (20 mM Tris-HCl pH 7.9, 25% glycerol, 420 mM NaCl, 1.5 mM MgCl2, 0.1% NP-40, 0.2 mM EDTA, 1 mM DTT, and 1 mM PMSF). After that, the nuclei were treated with Benzonase nuclease on ice for 30 mins and harvested by scraping. The soluble nuclear proteins were then separated from the insoluble chromatin fraction by centrifugation at 45,000 rpm for 1.5 h and then incubated with anti-FLAG beads (Thermo Fisher Scientific) with rotations overnight at 4°C. On the second day, after one wash with buffer 150 (20 mM Tris-HCl pH 7.9, 25% glycerol, 150 mM NaCl, 1.5 mM MgCl2, 0.1% NP-40, 0.2 mM EDTA, 1 mM DTT, and 1 mM PMSF), one wash with buffer 350 (20 mM Tris-HCl pH 7.9, 25% glycerol, 350 mM NaCl, 1.5 mM MgCl2, 0.1% NP-40, 0.2 mM EDTA, 1 mM DTT, and 1 mM PMSF), and two washes with buffer 150, the beads were eluted with reducing SDS-loading buffer, heated at 95°C for 10 mins, and analyzed by SDS–PAGE for subsequent Nano-HPLC-MS/MS analysis.

### Nano-HPLC-MS/MS analysis

Trypsin (12.5 ng/μl) was used for extract proteins in the gel slices. The peptides in the supernatant were collected after extract twice with solution (5% formic acid in 50% acetonitrile). Then, the tryptic peptides were subjected to NSI source, followed by tandem mass spectrometry (MS/MS) in Orbitrap Exploris 480 MS coupled online to the UPLC. After that, tandem mass spectra were extracted by Proteome Discoverer software (Thermo Fisher Scientific, version 3.0) and searched against Human database assuming the digestion enzyme trypsin. The acceptance criteria for identifications were the false discovery rate (FDR) should be less than 1% for peptides and proteins. Proteins identified in both replicate samples were subjected to downstream analysis.

### Total RNA-seq or RNA-seq

Total RNA was extracted as described above. Ribosomal RNA was removed by Ribo-off rRNA Depletion Kit (Human/Mouse/Rat) according to the manufacturés protocol (Vazyme) for total RNA-seq. RNA-seq experiments were performed using VAHTS Universal V6 RNA-seq Library Prep Kit for Illumina according to the manufacturés protocol (Vazyme). All total RNA-seq or RNA-seq libraries were performed with two biological replicates and sequenced on an Illumina NovaSeq 6000 platform.

### Chromatin immunoprecipitation (ChIP)

About 2 × 10^7^ Huh7 cells were cross-linked with 1% formaldehyde at room temperature for 10 mins, followed by quenching with 0.125 M glycine for 5 mins. Fixed cells were washed twice with ice cold PBS and resuspended in the cell lysis buffer (10 mM Tris pH 8, 10 mM NaCl, 0.2% Igepal) and nuclei lysis buffer (50 mM Tris pH 8, 1% SDS, 10 mM EDTA) with 1 × PMSF and 1 × protease inhibitors for 10 mins with rotations at 4°C. The nuclei were sonicated for 30 mins at 4°C (a train of 30 s on and 30 s off for 25-35 cycles). After removal of the insoluble debris, the rest of the supernatant was diluted with 3.5 ml IP dilution buffer (20 mM Tris-HCl pH 8.0, 2 mM EDTA, 150 mM NaCl, 1% Triton X-100 and 0.01% SDS) and pre-cleared with 50 μl Protein A/G agarose beads for 2 h at 4°C. After that, the supernatant was added Protein A/G beads pre-bound with 3-5 μg of specific antibodies against Flag-M2 (Sigma, A2220), SMARCA4 (Abcam, ab110641), H3K27ac (Abcam, ab4729), H3K4me3 (Abcam, ab8580), H2A.Z (Abcam, ab150402), H3K9me3 (Abcam, ab8898), H3K4me1 (CST, 5326), H3K27me3 (CST, 9733) and H4K20me3 (Abcam, ab9053) overnight. On the second day, after one wash with buffer I (20 mM Tris pH 8, 50 mM NaCl, 2 mM EDTA, 1% Triton X-100, 0.1% SDS), two washes with high salt buffer (20 mM Tris pH 8, 0.5 M NaCl, 2 mM EDTA, 1% Triton X-100, 0.01% SDS), one wash with buffer II (10 mM Tris pH 8, 0.25 M LiCl, 1 mM EDTA, 1% Igepal, 1% sodium deoxycholate), and two washes with TE (10 mM Tris pH 8, 1 mM EDTA pH 8), the beads were eluted twice with 100 μl elution buffer. And then, NaCl and RNase A were added to samples and incubated at 37°C for 30 mins. After that, proteinase K was added to samples and incubated at 65°C overnight. The DNA was purified with phenol-chloroform, followed by ethanol precipitation for library preparation.

### ATAC-seq

ATAC-seq experiments were performed using Hyperactive ATAC-Seq Library Prep Kit for Illumina according to the manufacturer’s instructions (Vazyme). All ATAC-seq libraries were sequenced on an Illumina NovaSeq 6000 platform. All ATAC-seq experiments were performed with two biological replicates.

### Single-cell RNA sequencing

Mouse HCC tissues were dissociated for single-cell RNA sequencing using the mouse tumor dissociation kit (130-096-730, Miltenyi Biotec) according to the manufacturer’s protocol. Briefly, the tissue was placed in a petri dish on ice and cut into small pieces of 2–4 mm, then infused with the RPMI/enzyme mix (Miltenyi Biotec), transferred to the gentleMACS Octo Dissociator, and run the program. After termination of the program, the suspension was passed through a 70 μm strainer and spun down at 300×g for 10 min at 4 °C, resuspended in PBS containing 0.04% BSA. Red blood cells were lysed with red blood cell lysis solution (130-094-183, Miltenyi Biotec) at 4 °C for 10 min, the resulting suspension was centrifuged at 300×g for 10 min and resuspended in PBS containing 0.04% BSA. Viability was assessed to be >85% using CountStar. Dissociated cells were resuspended in a final solution of PBS containing 0.04% BSA prior to loading on the 10× Chromium platform. The scRNA-seq libraries were generated using the 10× Genomics Chromium Controller Instrument and Chromium Single Cell 3’ V3 Reagent Kits (10× Genomics, Pleasanton, CA), according to the manufacturer’s recommendations. The experiments were performed by Berry Genomics Corporation (Beijing, China).

### Tatol RNA-seq or RNA-seq data processing and analysis

For Tatol RNA-seq or RNA-seq experiments, raw FASTQ files were aligned to the human (GRCh38/hg38) or mouse reference genome (GRCm38/mm10) using HISAT2 (65) with default parameters. SAMtools (66) was applied to transform SAM into the BAM format. FeatureCounts (67) was used to calculate the read counts of each transcript. TEtranscripts (68) was used to calculate the read counts of transposable elements. Analyzing repeat expression on repetitive elements was calculated by Homer (69) analyseRepeats.pl.

For differential expression analysis, we used *P* < 0.05 and fold-change > 2 as thresholds to identify differentially expressed genes by DESeq2 package (70). For differential transposable elements analysis, we used *P* < 0.05 and fold-change > 1 as thresholds. Commonly changed genes in both independent sgRNAs were considered to be significant. DAVID (71) was used for the GO and KEGG enrichment analysis. ClusterProfiler package (72) was used for Gene Set Enrichment Analysis. Data statistical tests were performed with GraphPad Prism.

### ChIP-seq data processing and analysis

For ChIP-seq experiments, raw FASTQ files were aligned to the human reference genome (GRCh38/hg38) using Bowtie 2 (73). SAMtools (66) was applied to transform SAM into the BAM format and removed PCR duplicates. Bigwig files were generated using deepTools (74) and the signal was normalized using the counts per million (CPM), followed by visualizing in the Integrative Genomics Viewer (IGV).

Peaks were generated by MACS2 (75) with the default parameters. The final peaks were those overlapped by both biological replicates, created by BEDTools (76). Peak annotation and sequence motif analysis were carried out with Homer (69). Heatmaps were generated by deeptools (74) computeMatrix and plotHeatmap.

### ATAC-seq data processing and analysis

For ATAC-seq experiments, raw FASTQ files were aligned to the human reference genome (GRCh38/hg38) using Bowtie 2 (73). All unmapped reads and PCR duplicates were removed by SAMtools (66). Bigwig files were generated and normalized using the CPM options in deepTools(74). MACS2(75) was used to call peak with the default parameters. Peak annotation and sequence motif analysis were carried out by Homer(69). Differential ATAC peaks were analyzed by DESeq2 package (70) with *P* < 0.05 and fold-change > 0.5 as thresholds. Heatmaps were generated by deeptools(74) computeMatrix and plotHeatmap.

### scRNA-seq data processing and analysis

For 10× Genomics Visium sequencing data, the raw reads were mapped to the mouse reference genome (refdata-gex-GRCm38-2020-A) using SpaceRanger software (v1.2.1, 10× Genomics). Briefly, Space Ranger showed the capture area of the organization in the chip by image processing algorithm and aligned reads of each spot according to the Spatial barcode information in Clean Data. Spot numbers, reads in spot, detected genes, and UMIs were counted. All spots detected by Space Ranger were included in the downstream analysis using R.

### Data availability

The RNA-seq, ChIP-seq, ATAC-seq and scRNA-seq data that support the findings of this study are deposited in the GEO repository (Accession Number: SRP15645607). and available from the corresponding author upon reasonable request. Data generated from the CRISPR genetic screen are provided in Supplementary Table S1.

### Statistical analysis

Results were presented as the mean ± SD. Different levels of statistical significance were accessed on the basis of specific *P* values with GraphPad Prism and R. *P* < 0.05 was represented as ‘*’, *P* < 0.01 was represented as ‘**’ and *P* < 0.001 was represented as ‘***’, respectively.

## Author contributions

X.L., J.C. and J.F. designed and conceived this project. R.G. initiated the project, performed most of the experiments, analyzed and interpreted data. R.G. and Y.S. conducted mouse studies. J.H. analyzed scRNA-seq patient data. J.W. provided TMAs and pathological analyses. X.S. assisted in analysis of the ChIP-seq and ATAC-seq data, and provided helpful discussions. J.H and S.L. analyzed TCGA patient data. Q. F., Y. J. and J. Z. helped with the experiments. X.L., J.C. and J.F. supervised the study. J. C. and X.L acquired funding. R.G., Y.S, J.H., J.W., J.F., J. C., and X.L. wrote the manuscript. All authors contributed to and approved the manuscript.

## Acknowledgments

We acknowledge the Medical Science Data Center at Shanghai Medical College of Fudan for providing the data analysis platform. This work was supported by the National Natural Science Foundation of China (32370614 to X. Lan). This work was also supported by grants from the National Natural Science Foundation of China (82272703 and 82473201 to J. Cai) and the Elite Youth Project of Natural Science Foundation of Fujian Province (2023J06056 to J. Cai).

