## Supplementary figures and images for "BRD8 Disruption Unlocks Retrotransposon-Driven Tumor-Intrinsic Immunogenicity via Redistribution of Histone Acetylation in Liver Cancer"

### Supplementary Fig. S1-S9

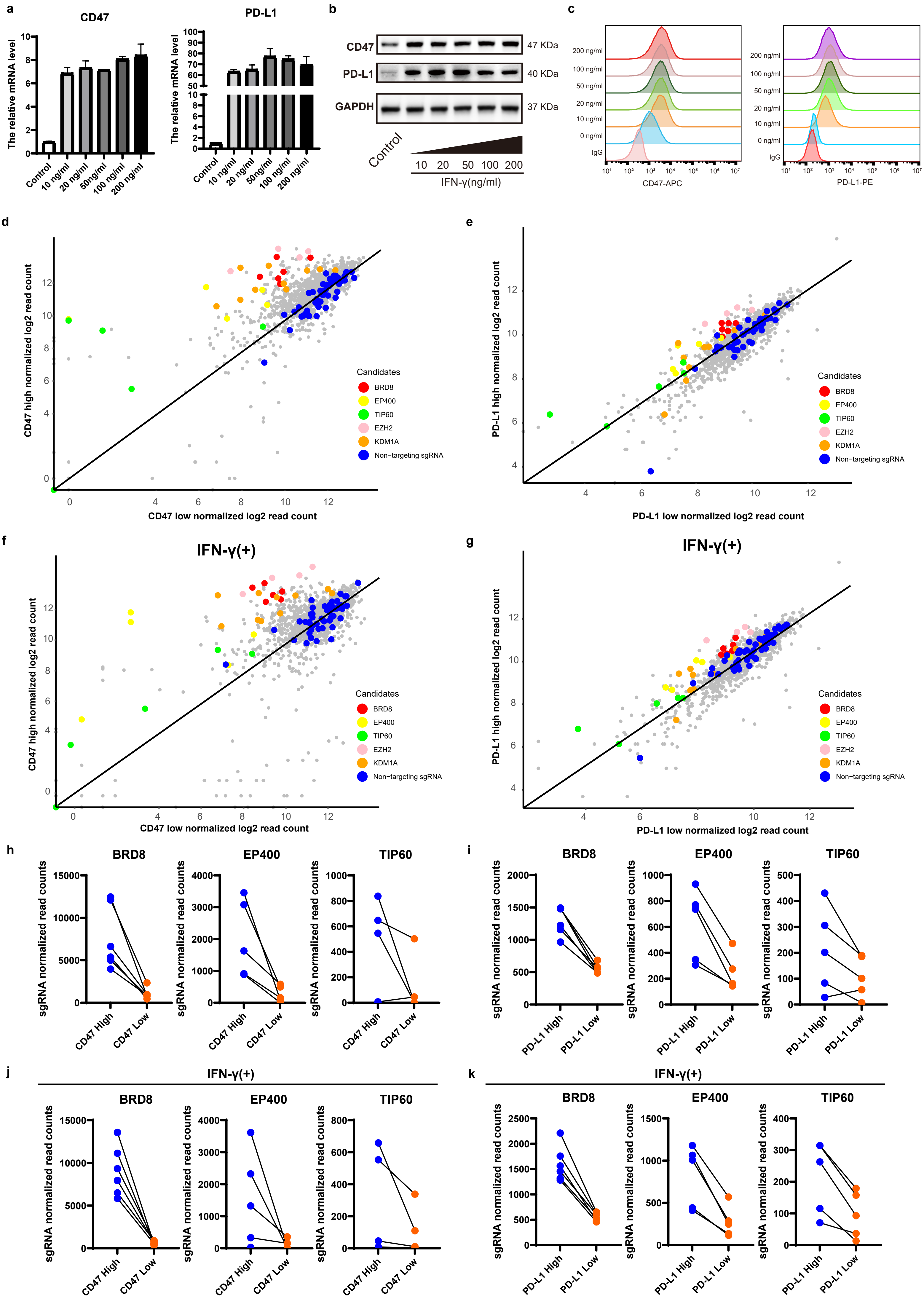

Supplementary Fig. S2

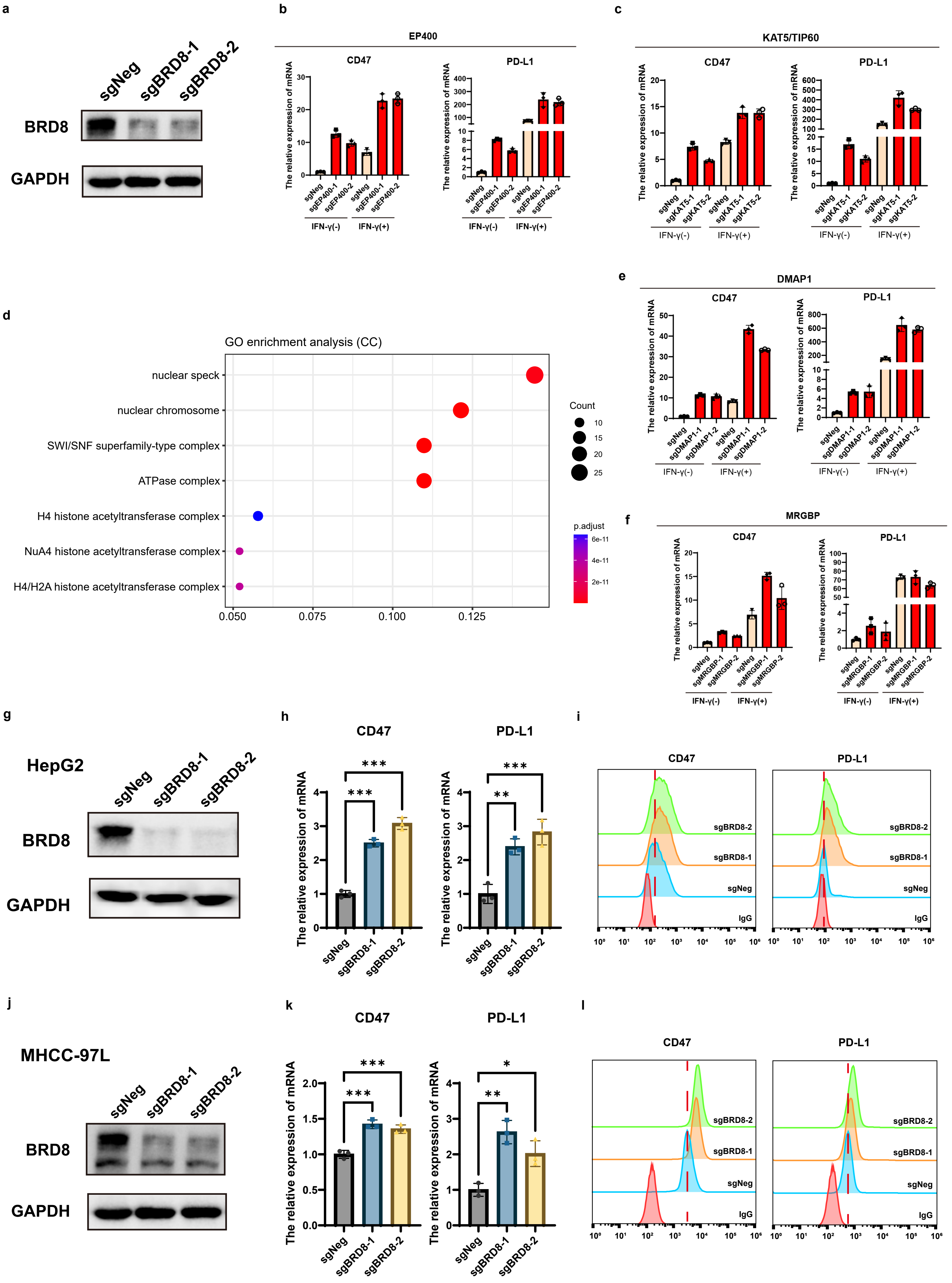

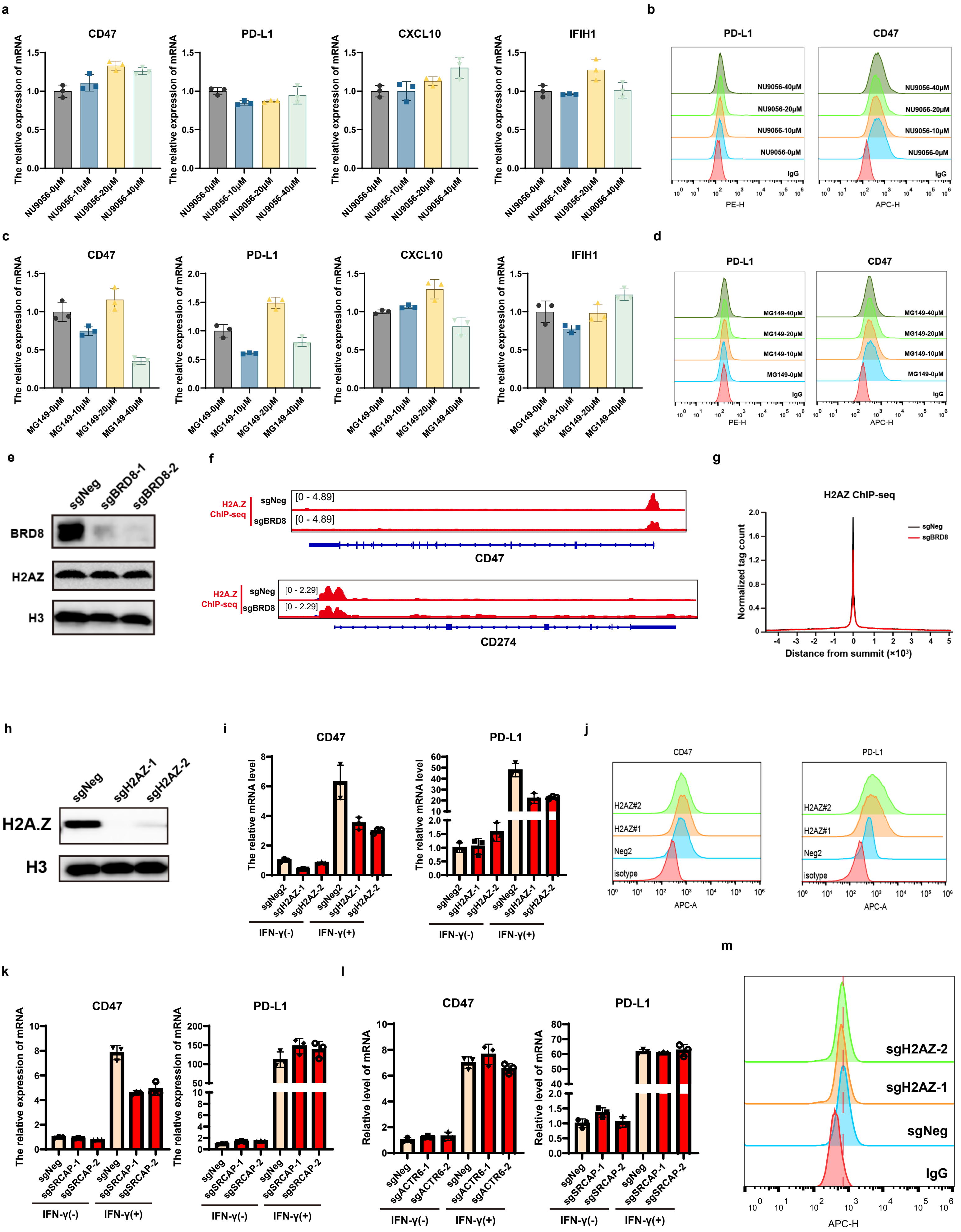

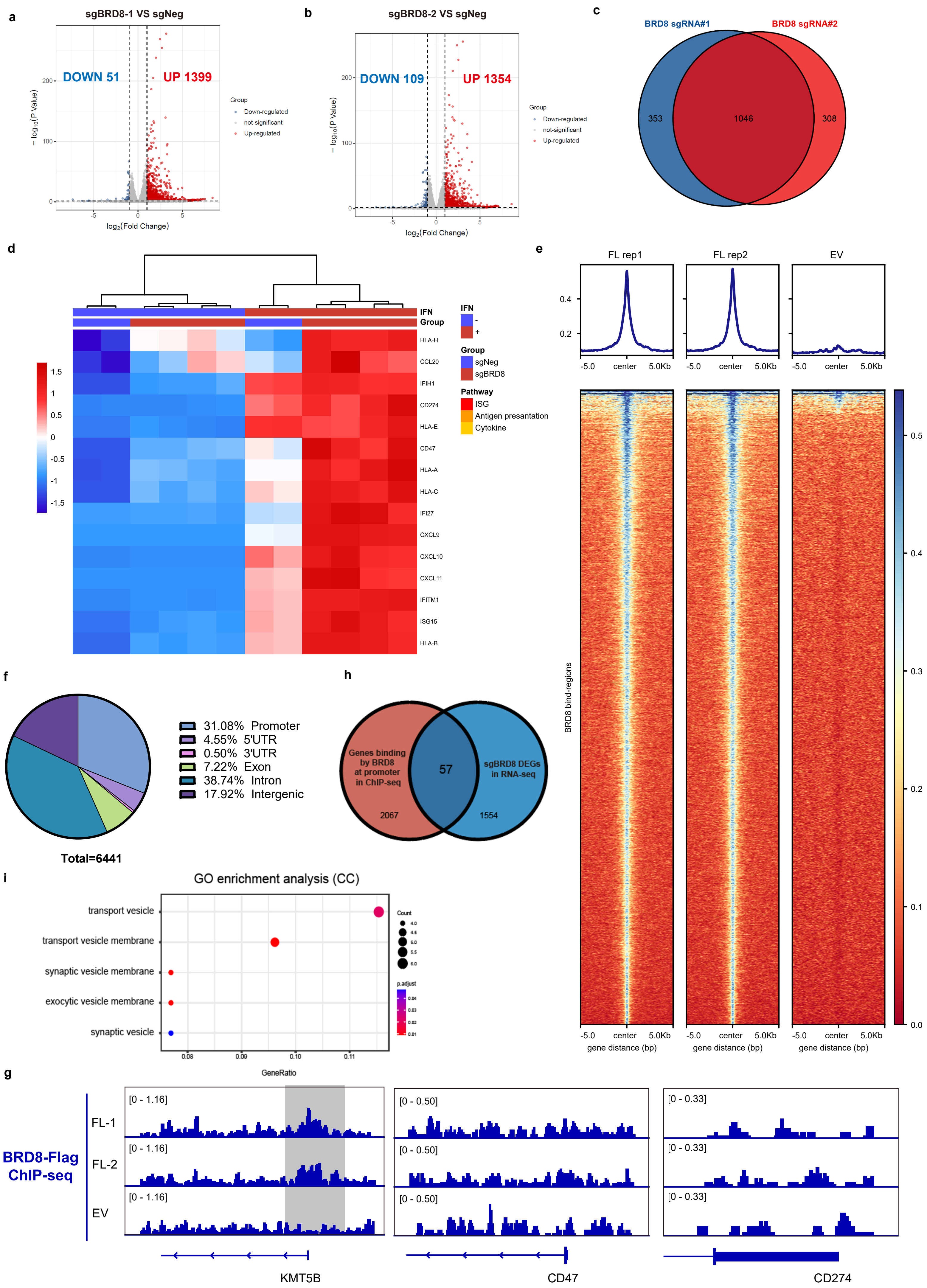

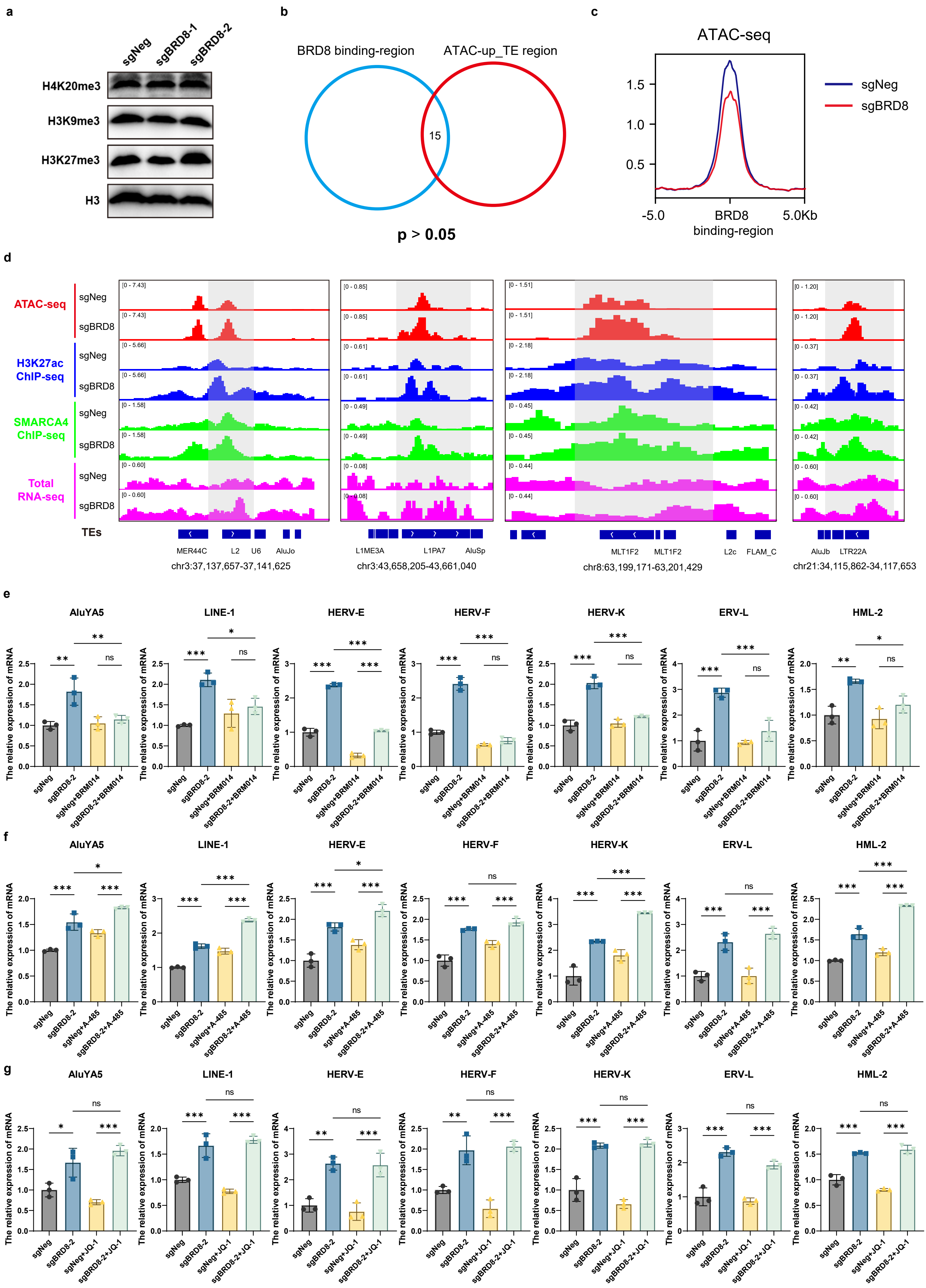

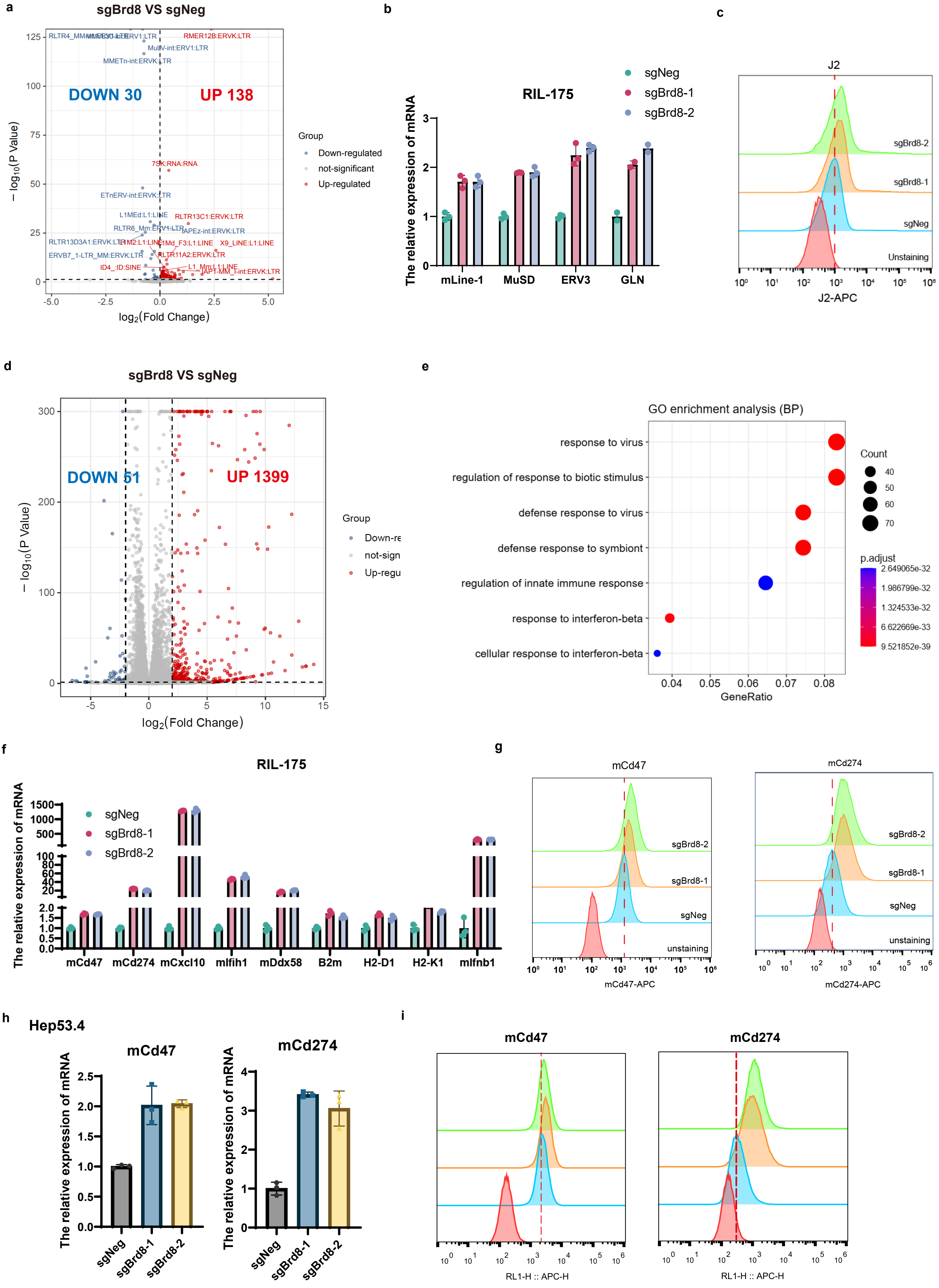

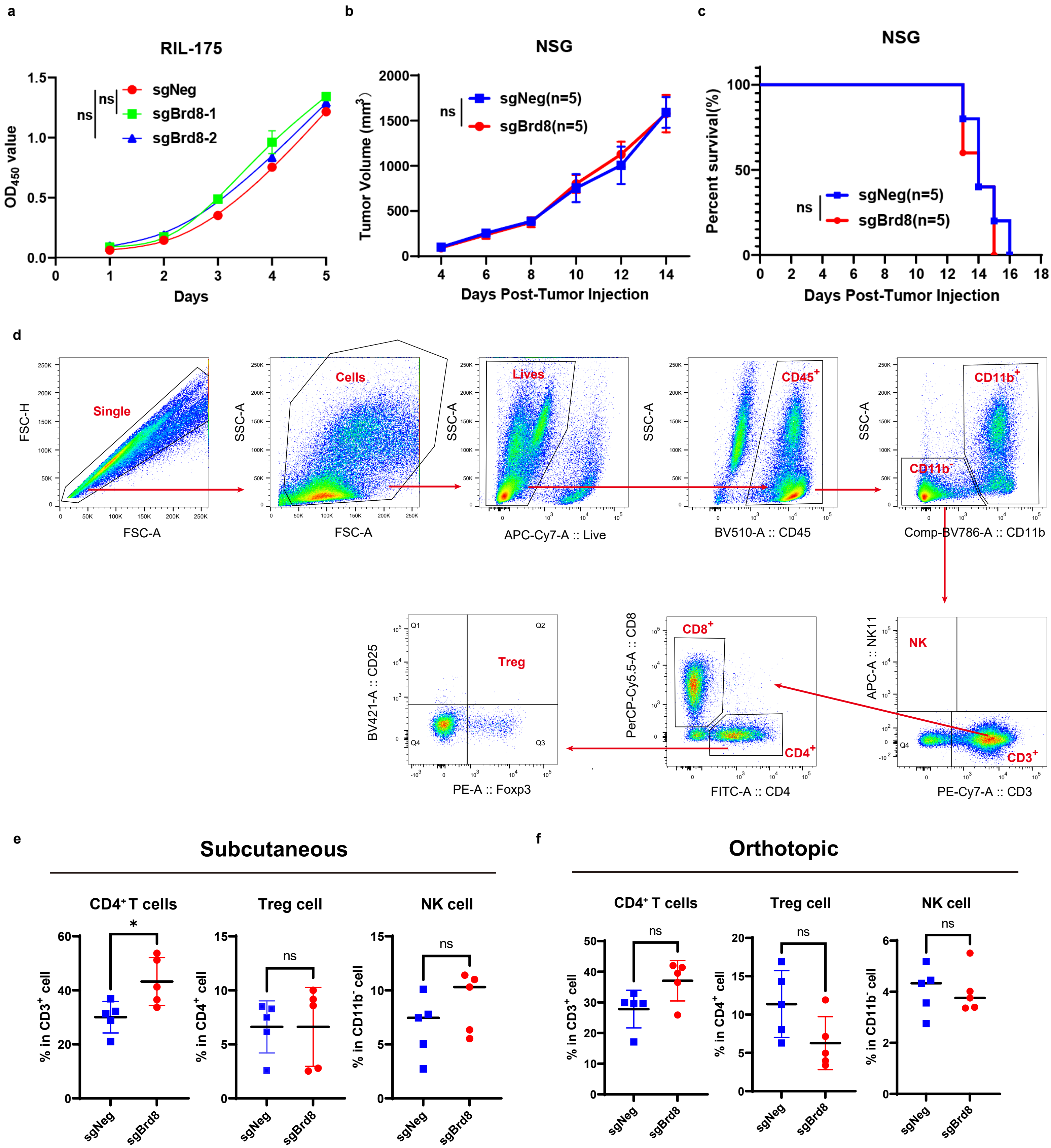

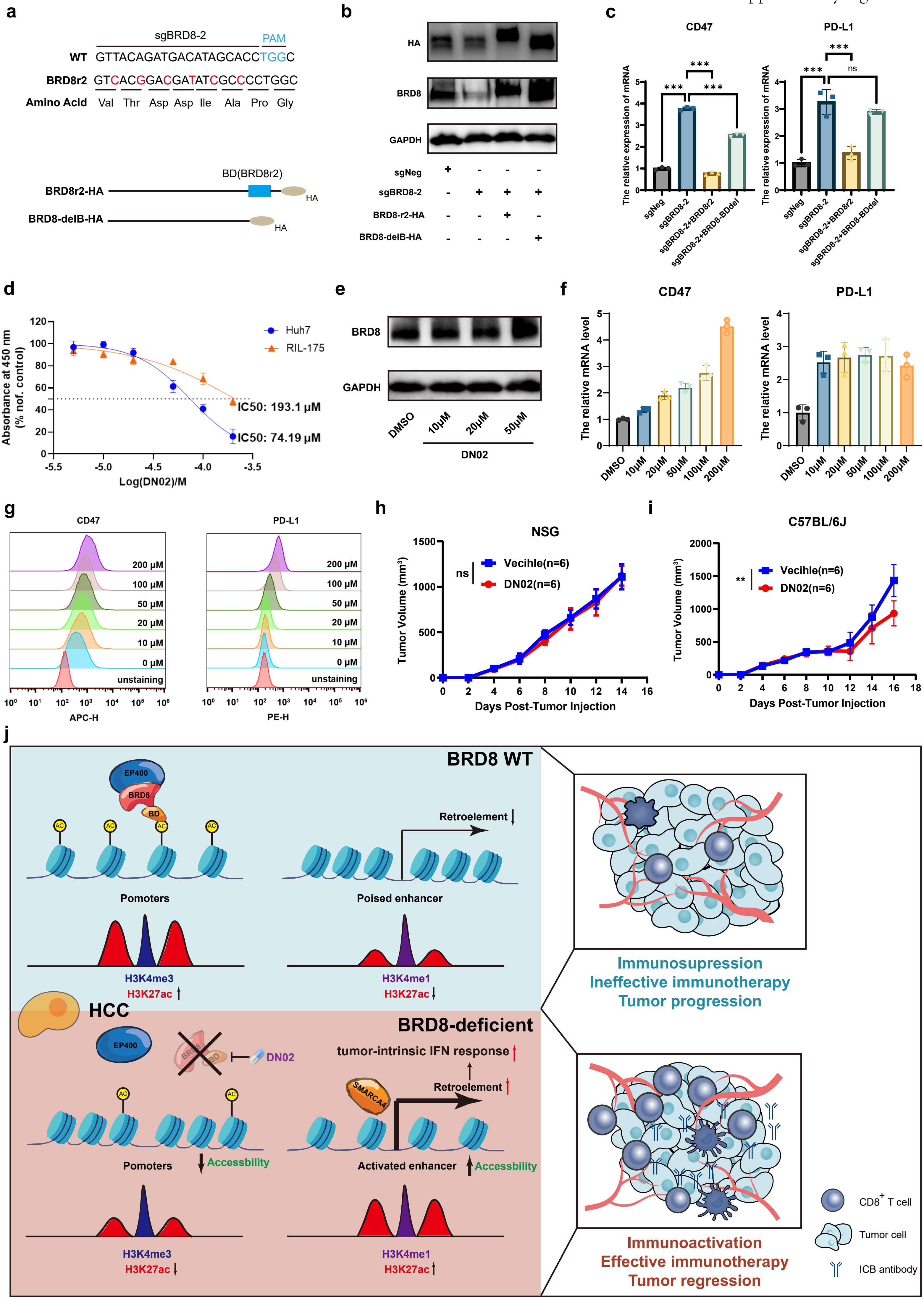

Supplementary Fig. S9

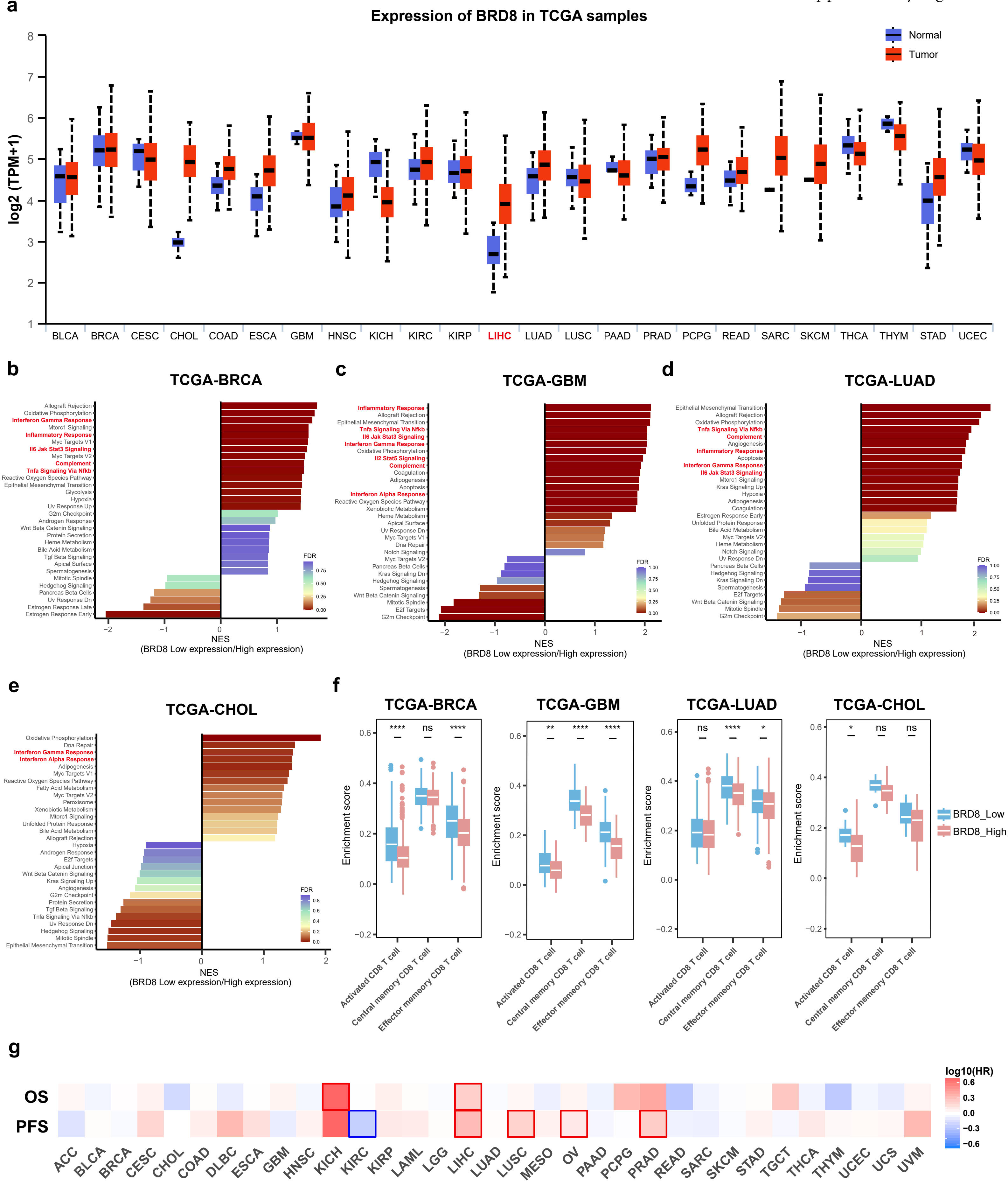
